# Allopregnanolone mediates affective switching through modulation of oscillatory states in the basolateral amygdala

**DOI:** 10.1101/2021.03.08.434156

**Authors:** Pantelis Antonoudiou, Phillip LW Colmers, Najah L Walton, Grant L Weiss, Anne C Smith, David P Nguyen, Mike Lewis, Michael C Quirk, Laverne C Melon, Jamie L Maguire

**Affiliations:** Department of Neuroscience, Tufts University School of Medicine, Boston, Massachusetts, 02111, USA; Sage Therapeutics, Inc., Cambridge, Massachusetts, 02142, USA; Department of Biology, Wesleyan University, Middletown, Connecticut, 06459, USA

## Abstract

Brexanolone (allopregnanolone), was recently approved by the FDA for the treatment of post-partum depression, demonstrating long-lasting antidepressant effects. Despite our understanding of the mechanism of action of neurosteroids as positive allosteric modulators (PAMs) of GABA_a_ receptors, we still do not fully understand how allopregnanolone exerts these persistent antidepressant effects. Here, we demonstrate that allopregnanolone and similar synthetic neuroactive steroid analogs, SGE-516 (tool-compound) and zuranolone (SAGE-217, investigational-compound), are capable of modulating oscillatory states across species, which we propose may contribute to long-lasting changes in behavioral states. We identified a critical role for interneurons in generating oscillations in the basolateral amygdala (BLA) and a role for delta-containing GABA_a_Rs in mediating the ability of neurosteroids to modulate network and behavioral states. Actions of allopregnanolone in the BLA is sufficient to alter behavioral states and enhance BLA high-theta oscillations (6-12Hz) through delta-containing GABA_a_ receptors, a mechanism distinct from other GABA_a_ PAMs, such as benzodiazepines. Moreover, treatment with the allopregnanolone analog SGE-516 induces long-lasting protection from chronic stress-induced disruption of network states, which correlates with improved behavioral outcomes. Our findings demonstrate a novel molecular and cellular mechanism mediating the well-established anxiolytic and antidepressant effects of neuroactive steroids.

## Introduction

New antidepressant treatments with proposed novel mechanisms of action, brexanolone and esketamine, recently received FDA approval for the treatment of postpartum- and treatment-resistant depression, respectively. Both of these treatments exert rapid and prolonged antidepressant effects which are not well explained by the proposed mechanisms of action of these compounds, offering the opportunity to explore potential novel mechanisms mediating these sustained antidepressant effects(Daly et al. 2018; Meltzer-Brody et al. 2018).

An interconnected network of brain areas including the basolateral amygdala (BLA) and prefrontal cortex (PFC) are critical for emotional processing in the brain(Calhoon and Tye 2015; Tovote, Fadok, and Lüthi 2015). Dysfunction of this network has been implicated in several neuropsychiatric disorders including depression, post-traumatic stress and anxiety(Babaev, Piletti Chatain, and Krueger-Burg 2018; Calhoon and Tye 2015; Fenster et al. 2018; Tovote, Fadok, and Lüthi 2015). Accumulating evidence suggests that rhythmic synchronization of BLA and PFC circuits is required for the behavioral expression of fear and anxiety(Davis et al. 2017; Felix-Ortiz et al. 2016; Karalis et al. 2016; Likhtik et al. 2014; Stujenske et al. 2014). Moreover, distinct oscillatory states in BLA/PFC areas seem to be associated with aversion and safety, the expression of which depends on inhibitory networks(Davis et al. 2017; Likhtik et al. 2014; Ozawa et al. 2020).

Here, we investigated the effect of neurosteroids on network activity in amygdalo-cortical regions implicated in mood and its effects on animal behavior. We show that allopregnanolone (and its analogs) alters oscillations in brain regions implicated in mood and promotes healthy network and behavioral states, providing a molecular and cellular mechanistic underpinning of neuroactive steroid-mediated affective switching(Schiller, Schmidt, and Rubinow 2014).

## Methods and Materials

Methods are described in detail in Supplement 1. EEG recordings were obtained from awake human cortical- and rat frontal-cortical regions. LFP recordings were obtained from the BLA of awake C57Bl/6J and global *Gabrd^-/-^* mice in response to administration of saline, allo (10mg/kg, i.p.), SGE-516 (5 mg/kg, i.p.), and diazepam (1 mg/kg, i.p.). Allopregnanolone-potentiated GABAergic currents were measured in BLA principal and PV+ interneurons using whole cell patch-clamp recording and cell type-specific expression of δ-containing GABA_a_Rs was examined using immunohistochemistry. Ex-vivo BLA oscillations were recorded in an interface recording chamber, induced by application of 800 nM kainic acid and elevated potassium (7.5 mM KCl). For ex-vivo optogenetic experiments, a viral vector (AAV8-EF1a-DIO-hChR2(H134R)-mCherry-WPRE-HGHpA) was delivered in the BLA of PV-cre mice. For optogenetic activation a blue light was delivered through a fiber optic (200 μm, 0.22 NA) coupled to a DPSS blue laser (473 nm, max power=500 mW, Laserglow technologies). For acute behavioral experiments, C57bl/6j and global *Gabrd-/-* mice were infused with saline solution (0.9 % NaCl) or 5 μg allo (2.5 μg/μl, Tocris). For chronic unpredictable stress (CUS), mice underwent a three-week protocol consisting of daily subjection to alternating stressors. The fMRI (blood-oxygen-level-dependent) BOLD signal was obtained using a Bruker BioSpec 7.0T with a 20-cm horizontal magnet and 20-G/cm magnetic field gradient quadrature transmit/receive coil (ID 38mm) at the Center for Translational Neuroimaging at Northeastern University.

## Results

### Allopregnanolone analogs alter brain oscillations across species

To examine whether SGE-516 and SAGE-217, with similar molecular pharmacology to brexanolone and allopregnanolone, alter brain states, we recorded cortical EEG from human subjects(Jobert et al. 2012) and rats. Oral application of SAGE-217 in healthy human subjects significantly elevated the power of the delta (δ), theta (θ) and beta1 (β1) frequency bands (δ: 42.7±12.30 %, n=7, p=0.01; θ: 33.9±9.70 %, n=7, p=0.01; β1: 34.3±11.80 %, n=7, p=0.03; unpaired t-tests Fig. 1 a-c). Therefore, SAGE-217 seems to produce robust alterations to human brain networks that could promote their anti-depressant effects and serve as a useful readout of target engagement.

**Figure 1.**
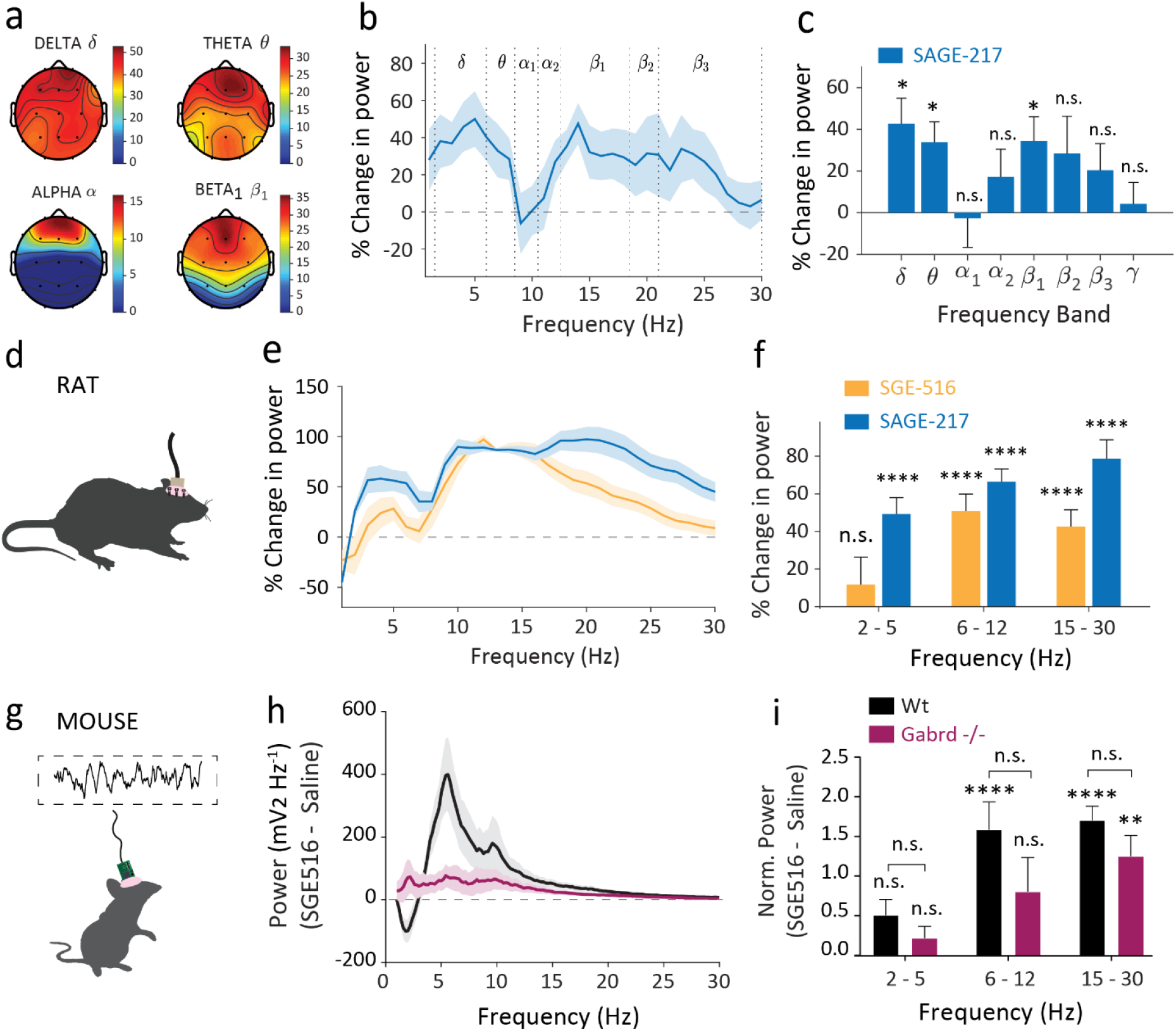
Neuroactive steroids altered brain network dynamics across species. **a-c, humans; d-f, rats; g-i, mice. a**, Heatmaps organized by frequency band showing regions where percentage change in Human EEG is elevated above vehicle. **b**, Power spectral mean (± SEM) difference from vehicle in EEG power (defined as percentage change from baseline) with SAGE-217 averaged from frontal electrodes F3, Fz, and F4. **c**, Mean (± SEM) change in power from vehicle (defined as percentage change from baseline) for SAGE-217. Shaded regions and error bars represent SEM (n=6 human subjects). **d**, Schematic for cortical EEG recordings in awake rats. **e**, Power spectral mean difference from vehicle in EEG power (defined as percentage change from baseline) in the rat with SGE-516 (orange) and SAGE-217 (blue). **f**, Mean (± SEM) change in power across frequency bands (n_SGE-516_=27 rats, n_SAGE-217_=20 rats); stars represent unpaired t-tests. **g**, Schematic for BLA LFP recordings in awake mice. **h**, Power spectral density difference between SGE-516 (5 mg/Kg) and saline IP applications. **i**, Normalized power area difference between acute SGE-516 and saline treatment across multiple frequency bands; Single stars represent paired t-tests between drug treatments. Brackets represent unpaired t-tests between genotypes (n_Wt_=6 mice, n_Gabrd_-/-=5 mice).

We also recorded the cortical EEG from rats treated with SGE-516 and SAGE-217 and quantified the power of oscillations in the low-theta (2-5Hz) band (associated with pro-fear states)(Davis et al. 2017; Karalis et al. 2016), high-theta (6-12Hz) band (associated with pro-safety states)(Davis et al. 2017) and beta band (15-30Hz). Similarly, both SGE-516 and SAGE-217 significantly increased power in high theta and beta bands in rats (SGE-516:: Norm. Power_SGE516-saline_^**6-12Hz**^: 50.71±9.13%, n=27, p<0.0001, unpaired t-test; Norm. Power_SGE516-saline_^**15-30Hz**^: 42.46±9.04%, n=27, p<0.0001, unpaired t-test; SAGE-217:: Norm. Power_SAGE217-saline_^**6-12Hz**^: 66.38±6.77%, n=20, p<0.0001, unpaired t-test; Norm. Power_SAGE217-saline_^**15-30Hz**^: 78.70±9.98%, n=20, p<0.0001, unpaired t-test; Fig. 1d-f). Additionally, SAGE-217 increased the power in the low theta band (Norm. Power_SGE516-saline_^**2-5Hz**^: 49.26±8.66%, n=20, p<0.0001, unpaired t-test; Fig. 1f). These results indicate that both allopregnanolone (allo) analogs elevate the power of oscillations consistently across high-theta and beta bands in rats.

Furthermore, we examined the effects of these analogs in mice on the LFP in the basolateral amygdala (BLA), an area implicated in the regulation of mood and anxiety(Babaev, Piletti Chatain, and Krueger-Burg 2018; Calhoon and Tye 2015; Tovote, Fadok, and Lüthi 2015). Acute IP injection of a non-sedative dose of SGE-516 (5 mg/kg) robustly increased the power in high theta and beta bands (Norm. Power_SGE516-saline_^**6-12Hz**^: 1.58±0.35, n=6, p<0.0001; Norm Power_SGE516-saline_^**15-30Hz**^: 1.70±0.18, n=6, p<0.0001, Sidak’s multiple comparisons; Fig 1g-i). On the other hand, acute IP injection of SGE-516 in mice lacking δ subunit GABA_a_Rs (Gabrd^-/-^; (Mihalek et al. 1999) altered only the beta band significantly compared to baseline (Norm. Power_SGE516-saline_^**15-30Hz**^: 1.25±0.26, n=5, p=0.0042; Norm. Power_SGE516-saline_^**6-12Hz**^: 0.80±0.43, n=5, p=0.06, Sidak’s multiple comparisons; Fig. 1h-i), indicating that the effects of SGE-516 on network states are partly mediated through GABA_a_R δ subunit-containing receptors. These findings suggest that the ability of allo analogs to alter oscillations in the theta and beta range are shared across species.

Benzodiazepine treatment is classically used to treat anxiety-related disorders. In contrast to neuroactive steroid GABA PAM treatment, diazepam (1 mg/kg) decreased the oscillatory power of the low theta band (Norm. Power_diazepam – saline_^**2-5Hz**^: −0.28±0.09, n=5, p=0.03, Sidak’s multiple comparisons; Supplementary Fig. 1a-c) and did not affect the oscillatory power of the high theta band (Norm. Power_diazepam - saline_^**6-12Hz**^: −0.10±0.06, n=5, p=0.43, Sidak’s multiple comparisons; Supplementary Fig. 1a-c) in BLA. These effects were not different between Wt and Gabrd^-/-^ mice (p^**2-5Hz**^>0.99, p^**6-12Hz**^=0.82, p^**15-30Hz**^=0.31, n_wt_=5, n_Gabrd_-/-=5, Sidak’s multiple comparisons; Supplementary Fig. 1c). In addition, a non-sedative dose of diazepam (2.5 mg/kg) did not affect the oscillatory power of cortical rat EEG in either the low or high theta bands (Norm. Power_diazepam - saline_^**2-5Hz**^: −16.66±19.78%, n=18, p=0.37, unpaired t-test; Norm. Power_diazepam - saline_^**6-12Hz**^: 9.23±11.04%, n=18, p=0.47, unpaired t-test; Supplementary Fig. 1d-f), again demonstrating similarity across species. In both mice and rats the beta-band power was enhanced after diazepam treatment (mouse: Norm. Power_diazepam - saline_^**15-30Hz**^: 0.29±0.029, n=5, p=0.0028, paired t-test; rat: Norm. Power_diazepam - saline_^**15-30Hz**^: 41.00±7.81%, n=18, p<0.0001, unpaired t-test; Supplementary Fig. 3c&f) in agreement with literature reports(Van Lier et al. 2004). These results suggest that neuroactive steroid GABA PAMs confer their effects through distinct network mechanisms from benzodiazepines.

### Allopregnanolone modulates BLA network oscillations in part through δ subunitcontaining GABA_a_ receptors

In order to investigate the effects of acute allo treatment on the BLA network, we recorded the LFP in the BLA of mice (Fig. 2a). Acute IP injection of allo (10mg/kg) produced robust changes in the pattern of BLA oscillations across multiple frequency bands. Specifically, acute allo administration strongly potentiated the normalized power in the high theta band (Norm. Power_allo - saline_^**6-12Hz**^: 0.83±0.117, n=13, p<0.0001, Sidak’s multiple comparison, Fig. 2b-e), signature oscillations associated with pro-safety states (Davis et al. 2017), and beta band (Norm. Power_allo - saline_^**15-30Hz**^: 1.18±0.117, n=13, p<0.0001, Sidak’s multiple comparison, Fig. 2e). To test whether these effects were mediated through GABA_a_R δ subunit-containing receptors, which are the predominant site of action for neurosteroids(Lee and Maguire 2014), we repeated the same experiments in Gabrd^-/-^ mice. Acute allo application also potentiated the normalized power in the high theta and beta bands (Norm. Power_allo - saline_^**6-12Hz**^: 0.20±0.054, n=5, p=0.02; Norm. Power_allo - saline_^**15-30Hz**^: 0.31±0.068, n=5, p=0.034, Sidak’s multiple comparisons, Fig. 2b-e) but to a significantly lesser extent than Wt mice (Abs. Power diff_wt,Gabrd-/-_^**6-12Hz**^: 0.64±0.194, p=0.0005, Abs. Power diff_wt,Gabrd-/-_^**15-30Hz**^: 0.86±0.196, p<0.0001, df=80, Sidak’s multiple comparisons). These findings suggest that acute allo treatment promotes the network communication of pro-safety oscillations in the BLA which are partially attributed to signaling via GABA_a_R δ subunit-containing receptors.

**Figure 2.**
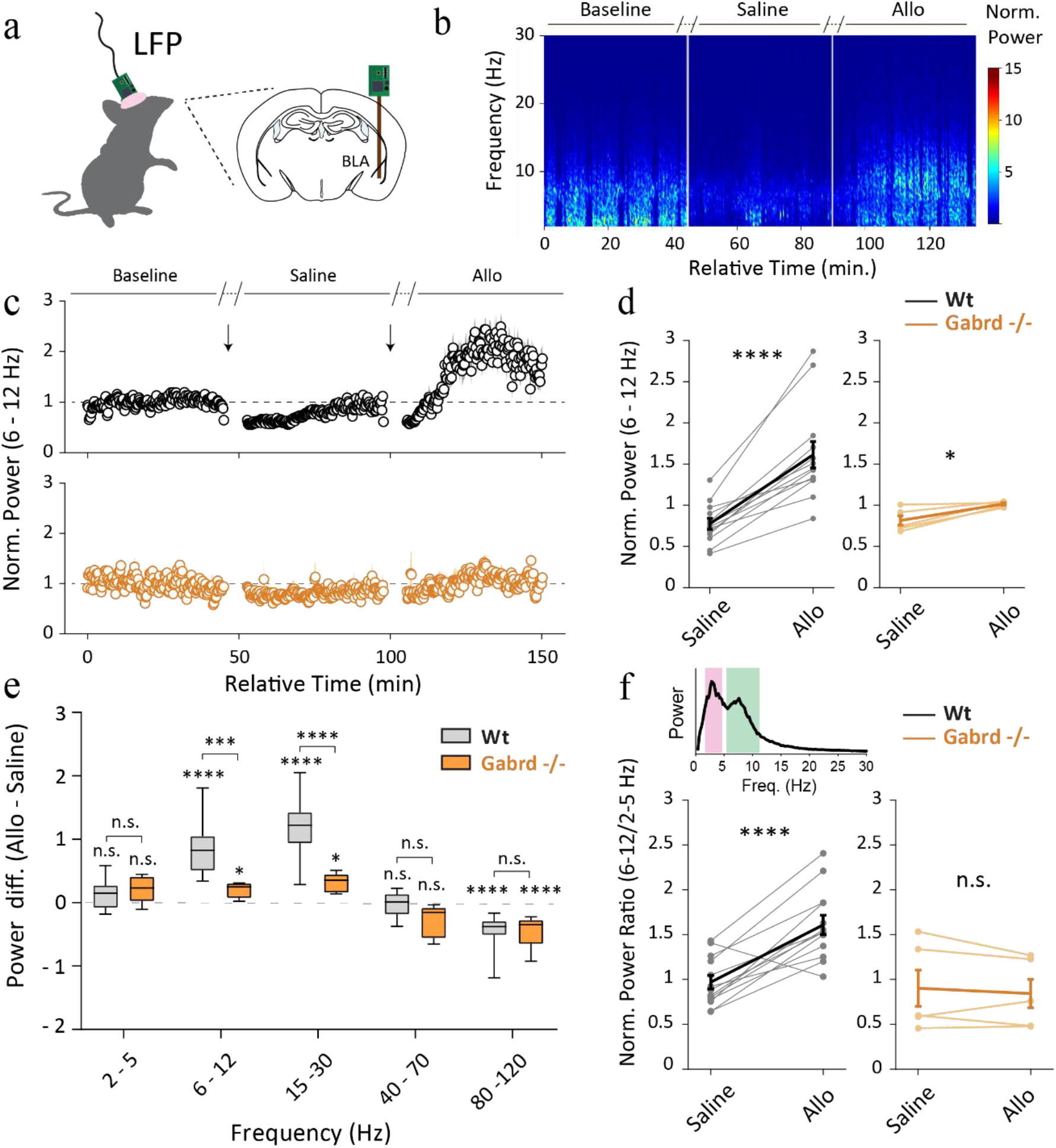
Acute IP application of allopregnanolone altered BLA network oscillations in freely moving mice partly through GABA_a_R δ subunit-containing receptors. **a**, Schematic for LFP recordings in awake mice. **b**, Representative spectrogram of BLA oscillations during IP application of saline and allo (10 mg/Kg). Normalized power^**6 - 12Hz**^ is higher during acute application of allo compared to saline treatment. **c**, Power area^**6 - 12Hz**^ normalized to baseline during saline and allo acute treatment across time **d**, Average dot plot; light color lines indicate individual experiments, dark lines the average and error bars the SEM. **e**, Normalized power area difference between acute allo and saline treatment across multiple frequency bands; Paired t-tests. **f**, High(green)/Low(pink) theta power ratio in Wt and Gabrd^-/-^ mice. n_Wt_=13 animals, n_Gabrd-/-_=5 animals.

On the other hand, acute allo application reduced the power of high-gamma^**80-120Hz**^ to a similar extent in both Wt and Gabrd^-/-^ mice (Wt:: Norm. Power_allo - saline_^**80-120Hz**^: −0.44±0.07, n=13, p<0.0001; Gabrd^-/-^:: Norm. Power_allo - saline_^**80 120Hz**^: −0. 44±0.12, n=5, p=0.0009, Sidak’s multiple comparisons; Fig. 2e) (Abs. Power diff_wt,Gabrd-/-_^**80-120Hz**^: −0.003±0.136, p>0.9999, df=80, Sidak’s multiple comparison). Therefore, allo reduced the power of BLA oscillations in the high-gamma band which have not previously been attributed to affective behavioral states and are independent of GABA_a_R δ subunit-containing receptors. The alterations observed in BLA oscillation power cannot be attributed to potentiation or habituation of the second IP injection as the LFP power to repeated saline injection is unchanged in both Wt and Gabrd^-/-^ mice (Supp. Figure 2).

To examine the relationship between fast and slow theta-band range oscillations, previously demonstrated to have opposing correlations with the behavioral expression of fear(Davis et al. 2017), we measured the power area ratio of high^**6-12Hz**^ and low^**2 - 5Hz**^ theta bands during IP application of saline and allo (10mg/kg). Acute allo application significantly increased the power ratio in comparison to saline injection in Wt mice, indicative of a shift to the signature oscillatory state associated with pro-safety(Davis et al. 2017), but not in Gabrd^-/-^ mice (wt:: Norm. Power_allo – saline_^**6-12/2-5Hz**^: 0.64±0.133, n=13, p<0.0001; Gabrd^-/-^:: Norm. Power_allo – saline_^**6 12/2 5Hz**^: −0.06±0.281, n=5, p=0.91, Sidak’s multiple comparison; Fig. 2f). Therefore, allo administration can switch BLA oscillations to the pro-safety state, an effect which seems to require GABA_a_R δ subunit-containing receptors.

### Allopregnanolone enhances tonic inhibition in BLA PV^+^ interneurons but not in principal cells

Our experiments suggest that allo actions in the BLA modulates network states through GABA_a_R δ subunit-containing receptors. Given the importance of PV^+^ interneurons in network coordination in BLA(Bartos, Vida, and Jonas 2007; Davis et al. 2017; Karalis et al. 2016; Veres, Nagy, and Hájos 2017) and the role of neuroactive steroids in potentiating tonic inhibition(Lee and Maguire 2014; Stell et al. 2003), we examined GABA_a_R δ subunit-containing receptor localization and the impact of allo on tonic inhibition on different cell types in BLA. We found that GABA_a_R δ subunit-containing receptors are uniquely expressed in PV^+^ interneurons in the BLA and are mostly absent in mice lacking the GABA_a_R δ subunit specifically in PV^+^ interneurons (PV-Gabrd^-/-^ mice) (Fig. 3a), as PV-Gabrd^-/-^ mice displayed a significant reduction in count for δ-positive cells in the basolateral amygdala when compared to Wt mice (wt: 17.00 ± 2.17; n=10 slices (3 mice), PV-Gabrd^-/-^: 4.09 ± 1.54; n=11 slices (4 mice); p<0.0001, Unpaired t-test; Fig. 3c). These data suggest that the expression of GABA_a_R δ subunit-containing receptors is largely restricted to PV^+^ interneurons in BLA.

**Figure 3.**
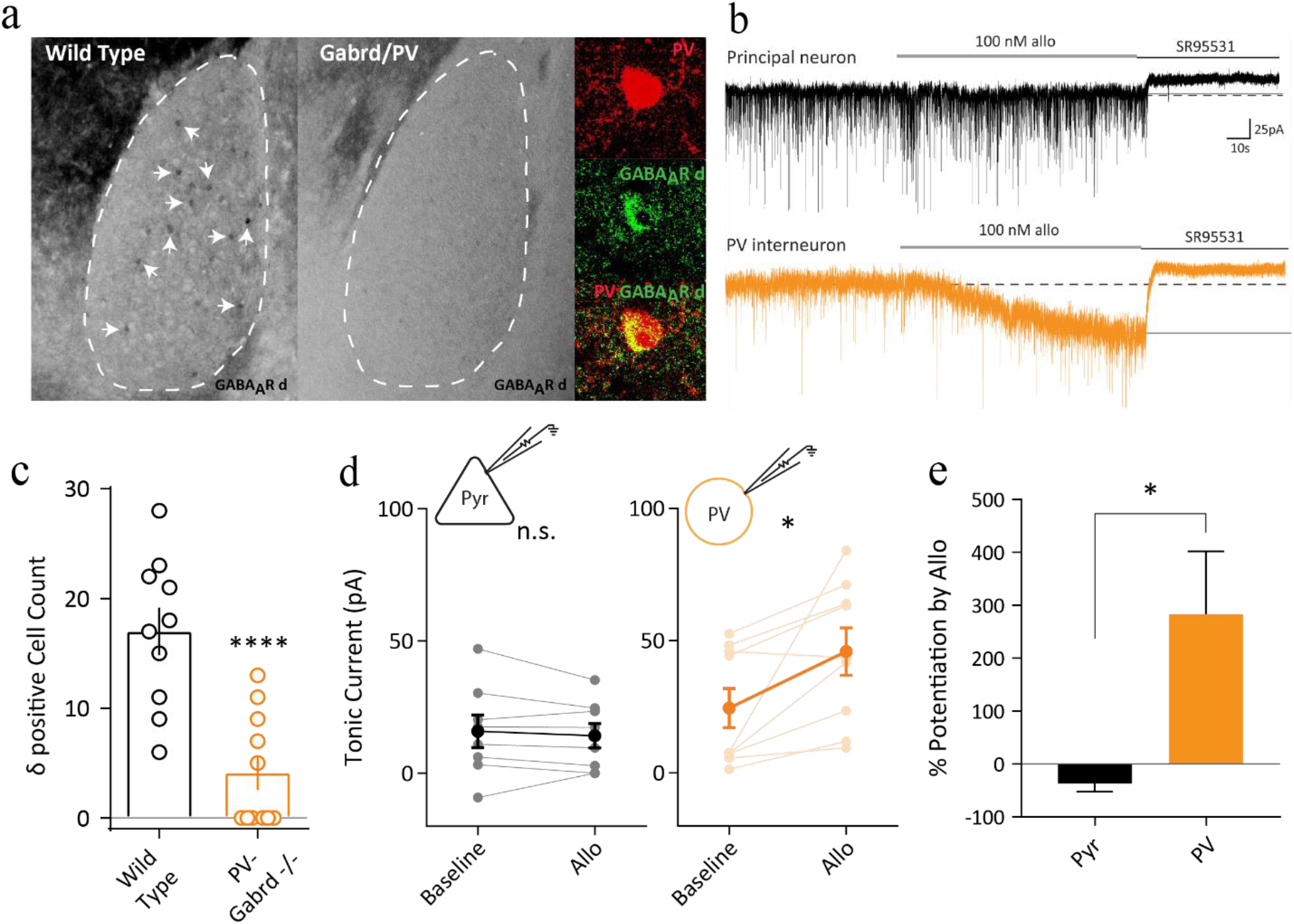
Allopregnanolone enhanced tonic currents in PV^+^ interneurons but not in principal cells of the BLA. **a**, PV^+^ interneurons highly express GABA_a_R δ subunit-containing receptors and immunoreactivity for these receptors is absent in PV-Gabrd^-/-^ mice. **b**, Representative traces from intracellular recordings in a principal and PV^+^ neurons during 100 nM allo application and block by SR95531 (Gabazine). **c**, PV-Gabrd^-/-^ mice displayed a significant reduction in δ immunoreactive cells in the basolateral amygdala when compared with controls (Unpaired t-test, t=4.921, wt: n=10 slices (3 mice), PV-Gabrd^-/-^: n=11 slices (4 mice)). **d**, Tonic current in principal cells (pyr) and PV^+^ interneurons (PV) during baseline and 100 nM allo application; paired t-test. **e**, Percentage potentiation of tonic current between allo and baseline application; Unpaired t-test with Welch’s correction. n_PV_=9 cells, n_Pyr_=10 cells.

To investigate the functional effects of GABA_a_R δ subunit-containing receptors, we performed whole cell patch-clamp recordings and applied 100 nM allo on BLA principal cells and PV^+^ interneurons, a concentration achieved during clinical trials with allo analogs, followed by SR95531 (≥100 μM) to unmask the tonic current. We found that allo strongly potentiated the tonic current in PV^+^ but not in principal cells (PV^+^: 21.34±7.68, n=9, p=0.02, paired t-test; Pyr: −1.66±2.19, n=8, p =0.47, paired t-test; Fig. 3b,d) resulting in a significantly higher potentiation of the tonic current between PV^+^ and principal cells (PV^+^= 282.9±118.7%, Pyr=-36.78±15.44; p=0.03; Unpaired t-test with Welch’s correction; Fig. 3e). These findings suggest that allo potentiates tonic currents in BLA PV^+^ interneurons through neurosteroid-sensitive GABA_a_R δ subunitcontaining receptors, which may contribute to the ability to orchestrate synchrony in the BLA (Pavlov et al. 2014).

### Isolated BLA networks can generate local brain oscillations

In order to examine the importance of PV^+^ interneurons in controlling the BLA circuit we sought to establish an *ex-vivo* oscillation model in isolated BLA networks of mice. Under regular aCSF conditions the LFP was silent (Supplementary Fig. 3a-b). However, addition of kainic acid (800 nM) and elevated KCl (High K^+^ / KA) into regular aCSF induced robust network oscillations at gamma (γ)-band frequency centered at 40Hz (Fig. 4a-c; Supplementary Fig. 3a-b). The current generator of these oscillations was situated within the BLA circuit, as nearby regions did not exhibit any oscillations (Supplementary Fig. 3c) and the hippocampus was removed to eliminate hippocampal volume conduction. Therefore, the isolated mouse BLA network is capable of intrinsically generating and self-sustaining local brain oscillations at gamma-band range^**30-80Hz**^. Furthermore, the generation of these oscillations does not seem to require phasic synaptic excitation as a selective AMPAR antagonist (Gyki 53655, n=7 slices; 6 at 20 μM, 1 at 10 μM) did not significantly alter gamma oscillation peak power (2.14±0.57 of baseline, one-sample t-test: p=0.09, n=7 slices (4 mice); Fig. 4d-e). On the other hand, gabazine application (10 μM) strongly suppressed gamma oscillation peak power (0.17±0.033 of baseline, one-sample t-test: p<0.0001, n=8 slices (4 mice); Fig. 4f-g) indicating that fast synaptic inhibition is critical for gamma oscillation emergence in BLA. Interestingly, following the collapse of gamma oscillations, rhythmic activity in the theta (θ) frequency range^**3-12Hz**^ emerged (368.52±165.61 of baseline, n=9 slices (4 mice); Fig. 4f-g). In 4 out of 9 of these experiments additional low-frequency events emerged (Supplementary Fig. 3e-f). Therefore, isolated BLA circuits are capable of generating network oscillations and fast-synaptic inhibition is critical for their emergence and may be critical in switching between different oscillatory states.

**Figure 4:**
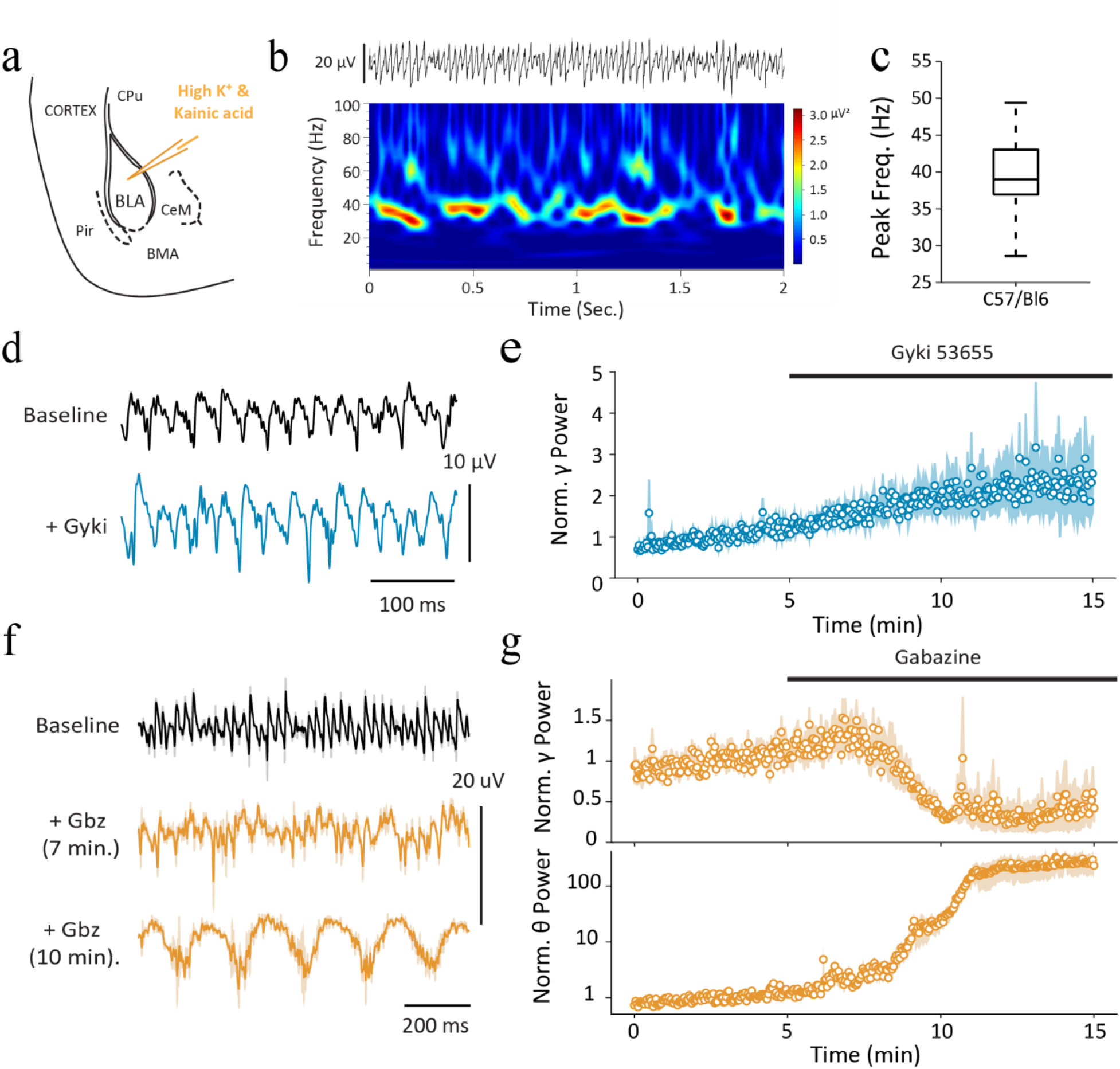
Isolated mouse BLA networks *ex-vivo* generated brain oscillations in the gamma and theta band-range. **a**, Schematic illustration of LFP setup for recording of *ex-vivo* gamma (γ) oscillations in mouse BLA slices. All experiments were performed in the presence of High K^+^/KA solution. **b**, top: Representative gamma oscillation LFP trace from BLA and bottom: Wavelet transformation. **c**, Peak frequency box plot of BLA gamma oscillations (n=42 slices). **d-e**, Gyki 53655 application to BLA gamma oscillations. **d**, Representative LFP traces during baseline (black) and Gyki 53655 application (blue). **e**, Normalized gamma peak power; dots represent mean and shaded region represents SEM (n=7 slices; 6 at 20 μM, 1 at 10 μM). **f-g**, Gabazine application to BLA gamma oscillations. **f**, Representative LFP traces during baseline (black) and gabazine application (orange). **g**, Normalized gamma (top) and theta (bottom) peak power change; dots represent mean and shaded region represents SEM (n=9 slices at 10 μM).

### PV^+^ interneurons can synchronize and control the BLA network

To assess whether fast-synaptic inhibition is sufficient to entrain the local BLA network and control oscillatory states we employed optogenetics to recruit BLA inhibitory interneurons. We targeted PV^+^ interneurons, which are the major class of GABAergic cells in BLA and have been suggested to provide powerful control over the somatic regions of local neurons(Bartos, Vida, and Jonas 2007; Krabbe, Gründemann, and Lüthi 2018; Veres, Nagy, and Hájos 2017). Infusion of AAV-DIO-ChR2-mCherry viral vector into the BLA of PV-cre mice resulted in expression of ChR2-mCherry in BLA PV^+^ interneurons (Fig. 5a-b). Pulsed blue light (473 nm) photo-excitation^**5-60Hz**^ produced robust phase-locked LFP responses suggesting that synchronized GABAergic release from PV^+^ interneurons can be detected in the BLA LFP (Fig. 5c-d; Supplementary Fig. 4a; p^**5Hz**^=0.0004, p^**10Hz**^<0.0001, p^**20Hz**^<0.0001, p^**30Hz**^=0.0001, p^**40Hz**^=0.0006, p^**50Hz**^=0.016, p^**60Hz**^>0.99; Bonferroni’s multiple comparisons test). In order to test if PV^+^ interneurons can entrain the BLA network and control BLA oscillatory state *ex-vivo* we examined the effects of pulsed opto-excitation during KA induced gamma. Pulsed blue light photo-excitation (5-60Hz) increased the power ratio (Fig. 5e-f; p^**5Hz**^=0.094, p^**10Hz**^<0.0001, p^**20Hz**^<0.0001, p^**30Hz**^<0.0001, p^**40Hz**^=0.076, p^**50Hz**^<0.0001, p^**60Hz**^=0.80; Bonferroni’s multiple comparisons test) and trains centered around 40Hz reliably entrained the ongoing kainate-induced gamma oscillations (Supplementary Fig. 4b-c). These results indicate that PV^+^ interneurons in BLA can powerfully control the local network synchronization across a range of frequencies. Furthermore, sustained photo-excitation of PV^+^ interneurons strongly suppressed ongoing gamma oscillations during light stimulation period (0.50±0.073 of baseline, n=23, p=0.0034, Dunnett’s multiple comparison; Fig. 5g-h) (Antonoudiou et al. 2020). This gamma power suppression was negatively correlated to light intensity (r=-0.42, p=0.0003, n=70, Spearman correlation, Fig. 5i), indicating that the stronger the light intensity the larger the gamma power suppression. Moreover, blue light stimulation had no effect in control slices from PV-cre mice that were not injected with the ChR2 viral vector (0.95±0.023 of baseline, n=6 slices, p=0.29, Dunnett’s multiple comparison; Supplementary Fig. 4d). Taken together these results suggest that GABAergic PV^+^ interneurons are powerful regulators of the local network activity and can control oscillatory states in BLA.

**Figure 5.**
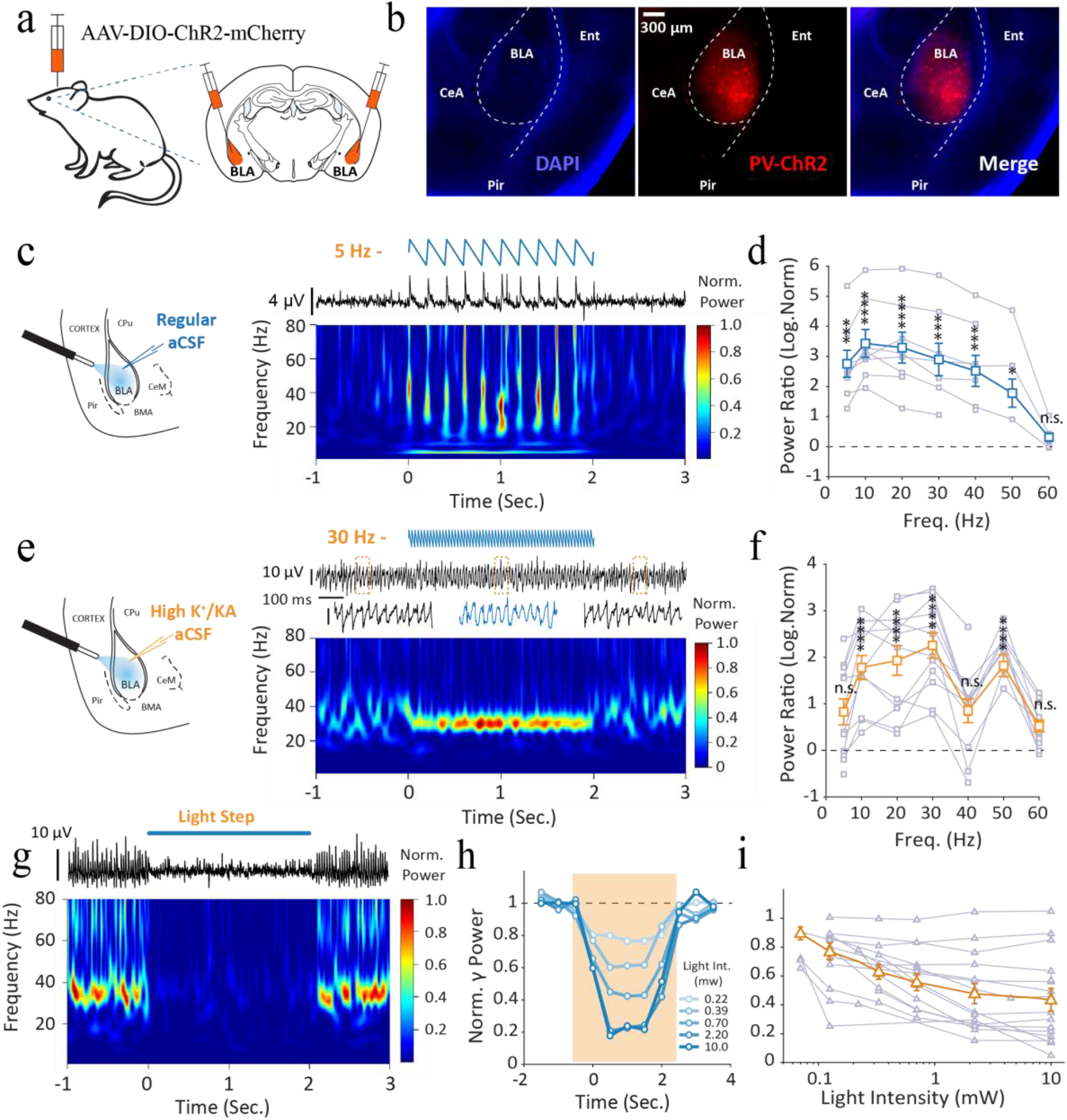
PV^+^ interneurons can synchronize and control the BLA network. **a**, Schematic of viral infusion of AAV-DIO-ChR2-mCherry in mouse BLA. **b**, Representative fluorescence images of ChR2-mCherry expression in BLA; CeA=Central Amygdala, BLA=basolateral amygdala, Pir=Piriform Cortex; Ent=Entorhinal Cortex. **c**, left-schematic illustration of LFP-opto setup for ChR2 experiments in aCSF; Right-top: Representative gamma oscillation LFP trace from BLA and bottom: Wavelet transformation. **d**, LFP power ratio between pulsed stimulation and baseline periods (n=8 slices). **e-f**, same as **c**-**d** during High K^+^/KA (n=12 slices) Bonferroni’s multiple comparisons test. **g-i**, Sustained PV^+^ interneuron excitation suppresses kainate-induced gamma oscillations. **g**, top: Representative gamma oscillation LFP trace from BLA and bottom: Wavelet transformation. **h**, Normalized gamma power to baseline across time for different light intensities. **i**, Gamma power normalized to baseline across light intensities (n=15 slices).

### Acute allopregnanolone infusion into the BLA alters behavioral states mediated through GABA_a_R δ subunit-containing receptors

In order to determine whether allo can exert effects in the BLA to alter avoidance behaviors, we implanted Wt mice with cannulas in the BLA (Supplementary Fig. 5a) and evaluated the impact of infusions on behavior. Allo infusion (5 μg) affected neither the total time spent (or their number of entries) in the open field center when compared with vehicle controls (Fig. 6a-c; time in Center_allo-veh_: 4.07±17.2 s, n_veh_=11, n_allo_=10, p=0.82, unpaired t-test; No. Entries-center_allo-veh_: 1.36±9.62, n_veh_=11, n_allo_=11, p=0.89, unpaired t-test), suggesting exploratory behaviors are unaltered by infusion of allo into the BLA. However, infusion of allo significantly increased the time that the mice spent in the lit area of the light dark box (time in lit area_allo-veh_: 75.17±35.03 s, n_veh_=11, n_allo_=13, p=0.043, unpaired t-test) and in the open arms of the elevated plus maze (time in open arm_allo-veh_: 61.24±22.05 s, n_veh_=11, n_allo_=12, p=0.011, unpaired t-test) when compared with vehicle controls (Fig. 6d-e & g-h). The number of entries made in the lit area of the light-dark box and in the open arm of the elevated plus maze did not differ between Wt mice that were infused with allo or vehicle (Fig. 6f & i; time in lit area_allo-veh_: 1.55±4.68, n_veh_=10, n_allo_=13, p=0.75, unpaired t-test; time in open arm_allo-veh_: −0.32±4.24, n_veh_=11, n_allo_=12, p=0.94, unpaired t-test). These results suggest that acute allo infusion into the BLA decreases avoidance behaviors in Wt mice, as indicated by the increased time they spend in anxiogenic regions of the behavioral apparatus. Further, mice treated with allo had a robust reduction in the amount of time spent immobile in the tail suspension test compared to vehicle treated controls (Fig. 6j-k; time immobile_allo-veh_: −79.29±11.15, n_veh_=12, n_allo_ =11, p<0.0001, unpaired t-test). These effects cannot be explained from impaired locomotor function as the total movement of vehicle and allo treated mice was not different in the open field and light dark box (Supplementary Fig. 5b-c; open-field_allo-veh_: −67.73±292.2 s, n_veh_=11, n_allo_ =11, p=0.82, unpaired t-test; time in lit area_allo-veh_ 105±227.4 s, n_veh_=11, n_allo_ =13, p=0.65, unpaired t-test). Importantly, these data demonstrate robust behavioral impacts of acute allo infusion restricted only on the BLA network.

**Figure 6.**
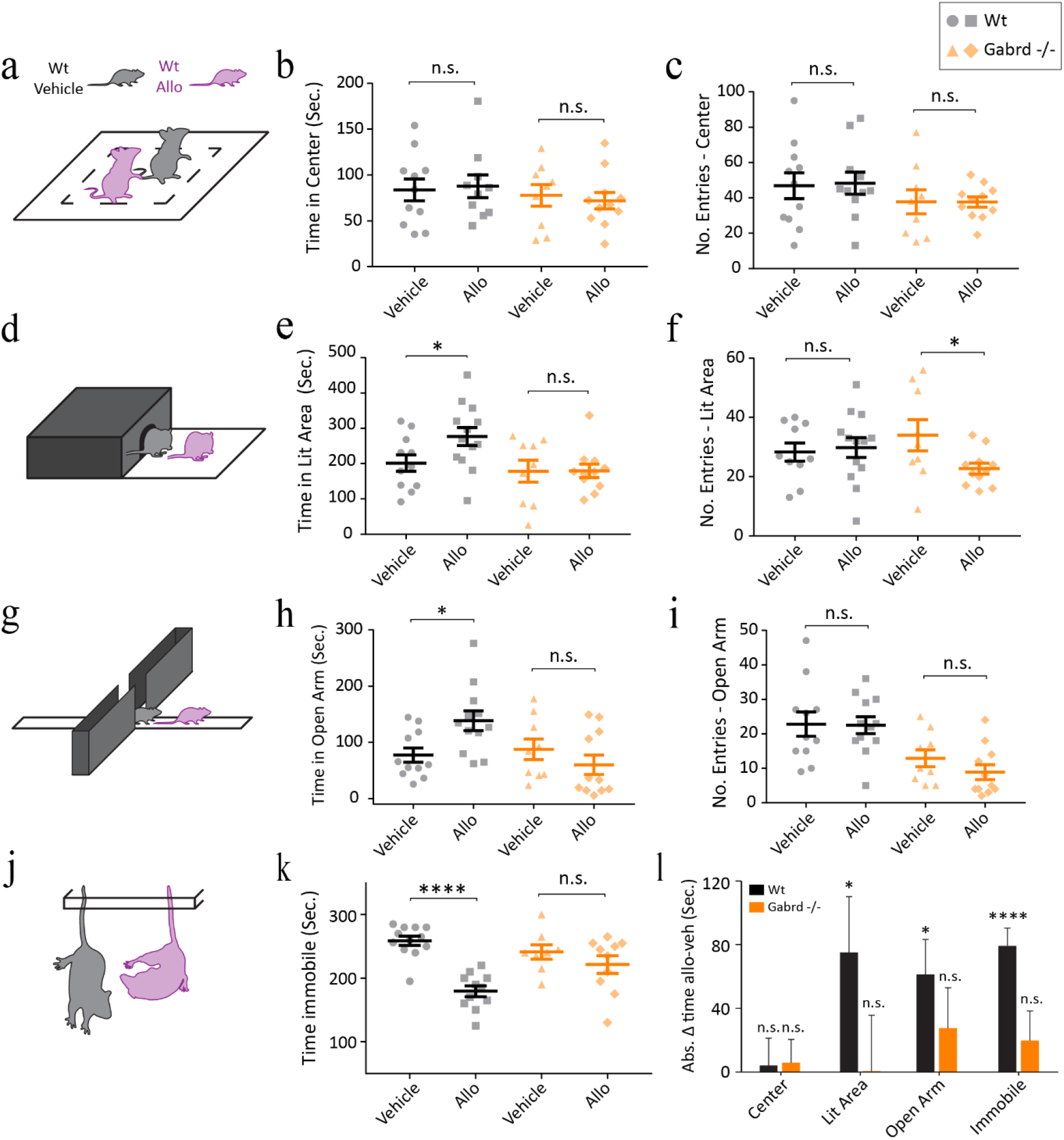
Allopregnanolone promoted anxiolysis in control but not in Gabrd^-/-^ mice. Schematics for behavioral tests and associated phenotypes in Wt mice infused with vehicle or 5 μg (2.5 μg/μl-inject) Allo: **a**, open field; **d**, light dark box; **g**, elevated plus maze, **j**, Tail suspension test. Time that animals spend in **b**, center; **e**, lit area; **h**, open arms, or **k**, immobile. Number of Entries in **c**, center; **f**, lit area; **i**, open arms. **l**, Summary of average absolute time difference in allo vs vehicle infused conditions across behavioral tests. Error bars represent SEM. Brackets and stars represent unpaired t-tests between vehicle vs allo treated groups.

To assess whether delta (δ) containing GABA receptors might play a role we repeated the same battery of test in Gabrd^-/-^ mice. Acute allo infusion in Gabrd^-/-^ mice (5 μg) did not change the time spent in the lit area of the light dark box (Fig. 6e; time in lit area_allo-veh_: 0.56±35.15 s, n_veh_=9, n_allo_ =11, p=0.99, unpaired t-test), open arm of the elevated plus maze (Fig. 6h; time in open arm_allo-veh_: −27.67±25.29 s, n_veh_=9, n_allo_ =11, p=0.29, unpaired t-test) or time immobile in the tail suspension test (Fig. 6k; time immobile_allo-veh_: −19.75±18.54 s, n_veh_=8, n_allo_ =10, p=0.30, unpaired t-test). Therefore, these data suggest that the behavioral effects of acute allo infusion into the BLA requires GABA_a_R δ subunit-containing receptors.

### SGE-516 prevents depressive-like behavior induced by chronic unpredictable stress

Our data show that neuroactive steroids alter behavioral states in WT mice. To assess the impact of neuroactive neurosteroids on sustained threat, WT mice were exposed to chronic unpredictable stress (CUS). Animals were subjected to 3 consecutive weeks of alternating stressors (see methods). LFP and behavioral tests were performed prior to and 4 weeks post CUS induction (Fig. 7a). The 4 weeks post CUS timepoint was chosen due to the demonstrated sustainment of allo efficacy at least 30 days post-injection in phase 3 clinical trials of post-partum depression(Meltzer-Brody et al. 2018).

**Figure 7.**
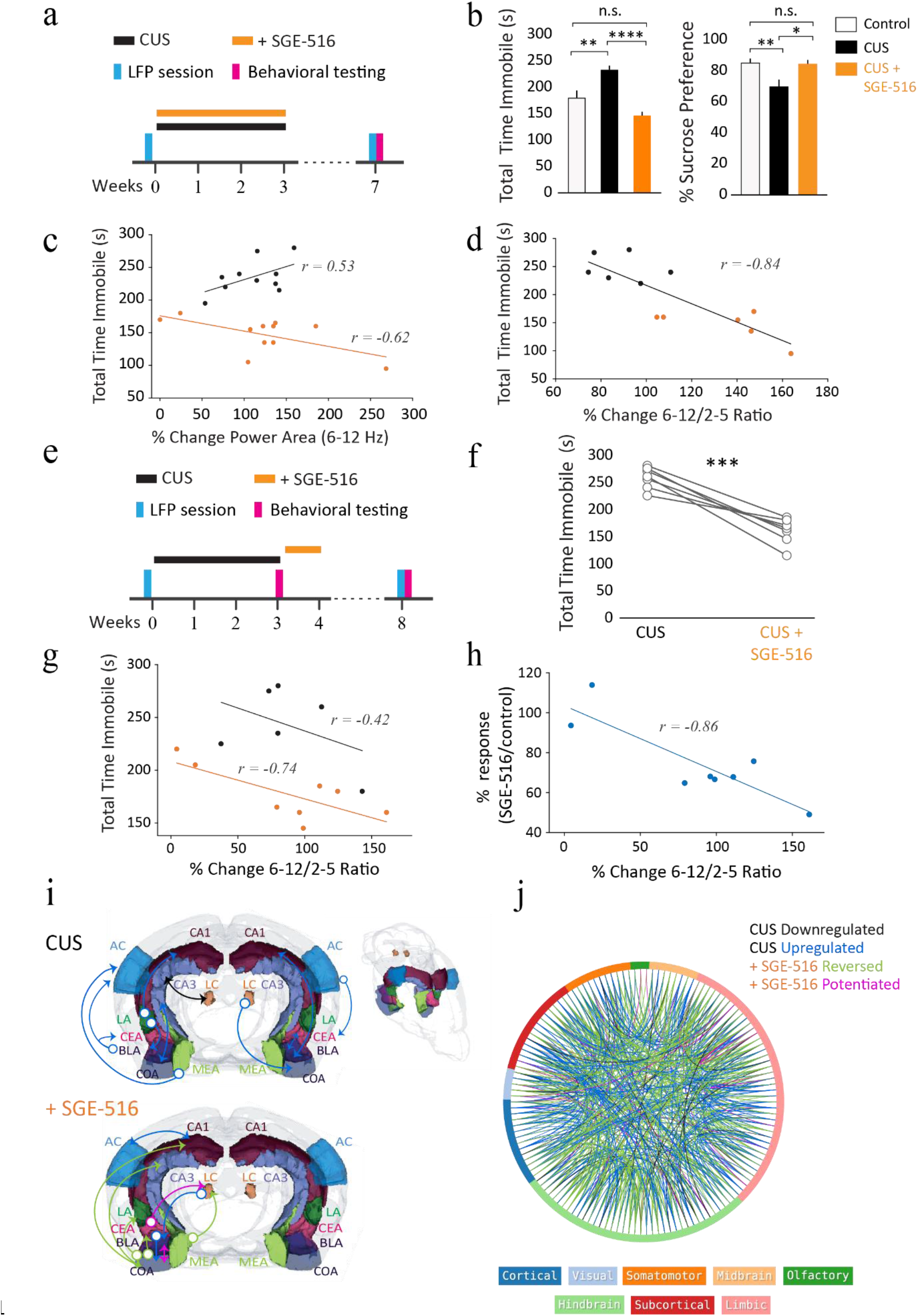
Synthetic Allopregnanolone analog SGE-516 prevents depressive-like behavior induced by chronic unpredictable stress (CUS) in C57bl/6 mice. **a-d**, SGE-516 application during CUS. **a**, Timeline of CUS paradigm with LFP and behavioral sessions. **b**, Left - Time spent immobile in tail suspension test (TST) and Right-% sucrose preference in control/no CUS (white), CUS (black), and CUS +SGE-516 (orange) animals. **c**, Time spent immobile in TST plotted against high theta (6-12Hz) power change prior to and post CUS induction in CUS (black) and CUS + SGE-516 (orange) mice. **d**, Time spent immobile in TST plotted against theta ratio change (6-12/2-5HzHz) in both CUS (black) and CUS +SGE-516 (orange) using outlier selection. **e-h**, SGE-516 application after CUS. **e**, Timeline of CUS paradigm with LFP and behavioral sessions. **f**, Time spent immobile in TST before and after SGE-516 application. **g**, Time spent immobile in TST plotted against theta ratio (6-12/2-5HzHz) change prior to and post CUS induction in CUS (black) and CUS + SGE-516 (orange) mice. **h**, TST response difference prior to and post SGE-516 application plotted against theta ratio (6-12/2-5HzHz) change prior to and post CUS. Power changes in correlation plots were calculated as the percent difference between 4 weeks post-CUS and 24 hours prior to CUS; *r*-values represent the Pearson correlation coefficient. **i-j**, Resting state functional connectivity; ***type-of-connection legend:*** black: downregulated, blue: upregulated, green: reversed by SGE-516, pink: potentiated by SGE-516. **i**, Representative brain volumes of resting state functional connectivity in CUS and CUS+SGE-516 mice. Arrows indicate directionality of connections from the indicated seed regions; arrow color reflects type of connection change based on type-of-connection legend; *AC: Auditory Cortex, BLA: Basal Lateral Amygdaloid Area, CA1: Cornu Ammonis 1, CA3: Cornu Ammonis 3, CEA: Central Nucleus of the Amygdala, COA: Cortical Amygdaloid Area, LA: Lateral Amygdaloid Area, LC: Locus Coeruleus, MEA: Medial Amygdaloid Area.* **j**, Connectome map indicating significant changes in BOLD signal connectivity, Outer circle lines represent brain areas of the respective color as indicated in the bottom legend.

Mice that underwent CUS spend more time immobile in the tail suspension test (Fig. 7b; Control: 180.0±13.76 s, n=10 animals; CUS: 233.2±7.75 s, n=11 animals; p=0.0019, Tukey’s multiple comparison test) and had a lower preference for sucrose than control mice (Fig. 7b; Control: 85.3±2.72%, n=15 animals; CUS: 70.0±4.41%, n=10 animals; p=0.0053, Tukey’s multiple comparison test). SAGE-516 application during CUS had the opposite effects as it decreased time immobile in the tail suspension test (Fig. 7b; CUS + SAGE-516: 146.3±7.49 s, n=12 animals, p<0.0001, Tukey’s multiple comparison test) and elevated sucrose preference (Fig. 7b; CUS + SAGE-516: 84.6±2.56%, n=8, p=0.0243, Tukey’s multiple comparison test) near to levels observed in controls (Fig. 7b). These results suggest that sustained threat through CUS induces behavioral deficits in WT mice which are prevented by SGE-516 treatment. To investigate how BLA network activity is associated with behavioral changes, we compared BLA high theta (6-12Hz) power change prior to and post CUS induction with time spend immobile in the tail suspension test. We found that mice that underwent CUS had a positive non-significant correlation (Fig. 7c; *r* = 0.53, n=10 animals, p=0.11, Pearson correlation) between time immobile and high theta change. However, there was a significant negative correlation between (Fig. 7c; *r* = −0.62, n=11 animals, p=0.041, Pearson correlation) time immobile and high theta change in CUS mice that were treated with SGE-516, indicating that larger increases in high theta power were associated with less time spend immobile and depressive-like behavior in mice (z = 2.54, p = 0.01, Fisher transformation). Furthermore, when including animals with the most extreme changes in behavioral outcomes from both CUS and CUS+SAGE-516 groups, there was a strong negative correlation (Fig. 7d; *r* = −0.84, n=12 animals, p=0.00057, Pearson correlation) between time spent immobile and change of high/low (6-12/2-5Hz) theta power. These data suggest that larger high/low theta power increases in BLA are associated with improved behavioral outcomes.

Our results show that SGE-516 can counteract the behavioral deficits induced by sustained threat. In order to see if they can also reverse these behavioral deficits, we treated a separate group of animals with SGE-516 after CUS induction (Fig. 7e). SGE-516 treatment following CUS exposure robustly reduced the total time spent immobile during a tail suspension test tests 4 weeks following treatment (Fig. 7f; CUS: 257.9±7.47 s, CUS + SGE-516: 160±9.00, n=7, p=0.0003, paired t-test). Therefore, our data suggest that treatment with allo analog SGE-516 reduces behavioral deficits induced by sustained threat. Moreover, in CUS mice treated with SGE-516 increases in high/low theta power from prior to post-CUS were associated with reduced time spent immobile in the tail suspension test (Fig. 7g; *r* = −0.74, n=8 animals, p=0.035, Pearson correlation). In addition, increased high/low theta ratio change in BLA (post-CUS/pre-CUS) was associated with the behavioral response elicited from SGE-516 treatment (Fig. 7h; *r* = −0.86, n=8 animals, p=0.006, Pearson correlation). Overall, these results suggest that increases in high/low theta ratio are associated with a reduction in behavioral deficits resulting from sustained threat in mice.

To more broadly and unbiasedly examine the effects of CUS on global brain connectivity we acquired the fMRI blood-oxygen-level-dependent (BOLD) signal. Mice that underwent the CUS protocol exhibited altered functional connectivity across multiple brain areas when compared to controls (Fig. 8i-j; CUS altered 84.95% of control connections: 254/299 and 325 new connections). Interestingly, SGE-516 treatment during CUS reversed 47.32% (274/579) of the CUS altered connections restoring control connectivity (Fig. 8j). Additionally, 89.78% (246/274) of the reversed network connections were baseline control connections, indicating most of the new connections made by CUS were not affected by SGE-516 treatment. These data show that sustained threat produces brain-wide changes in functional connectivity of WT mice and suggest that treatment with a synthetic neurosteroid analog of allo could act to partially restore those changes.

## Discussion

Here we demonstrate that neurosteroids alter network oscillations across multiple species (Fig.1) and in rodents they potentiate high-theta^**6-12Hz**^ power unlike other GABA agonists (Fig.1-2 & Supplementary Fig.1). This increase in BLA high-theta power was found to be largely mediated through δ subunit-containing GABA_a_Rs which are uniquely expressed in PV+ interneurons in the BLA (Fig. 2-3). Furthermore, the BLA network *ex-vivo* can self-sustain oscillations that are under PV^+^ interneuron control (Fig.4-5). Importantly, we show that acute allo infusion into the BLA is sufficient to alter behavioral states (Fig. 6) and systemic neurosteroid (SGE-516) treatment protects mice from behavioral deficits induced from sustained threat (Fig. 7).

It has been shown that distinct oscillatory states in BLA-PFC areas seem to be associated with aversion and safety(Davis et al. 2017; Karalis et al. 2016; Likhtik et al. 2014). Indeed, the behavioral expression of fear and safety associated with shifts in the low-^**~4Hz**^ and high-theta^**~8Hz**^ bands, respectively(Davis et al. 2017; Karalis et al. 2016; Ozawa et al. 2020). Here, we observed that acute allo increased the high/low theta^**6-12/2-5Hz**^ power in the BLA (Fig. 2f) and decreased behaviors relevant to anxiety- and depressive-like states (Fig. 6). Furthermore, allo-analog (SGE-516) treatment restored healthy behavioral states in mice that underwent sustained threat (Fig. 7e-f), where larger restoration was associated with greater increases in BLA high/low theta^**6-12/2-5Hz**^ power (Fig. 7g-h). Therefore, our findings reinforce the evidence that increases in high/low theta band are associated with safety(Davis et al. 2017; Ozawa et al. 2020) and further suggest that they could generalize them to healthy brain states beyond the realm of fear. We also observed that high-theta^**6-12Hz**^ and beta^**15-30Hz**^ oscillations in BLA were significantly higher during allo application in WT compared to *Gabrd^-/-^* mice (Fig. 2e) indicating that they could also be an important component of neurosteroid treatment. Beta^**15-30Hz**^ power was also strongly elevated by diazepam in both mice and rats (Supplementary Fig. 1), in agreement to previous findings(Van Lier et al. 2004). These results could suggest that high-theta^**6-12Hz**^ and not beta^**15-30Hz**^-oscillation elevation is unique to the proposed mechanisms of action mediated by neuroactive steroids and potentially contribute to their superior antidepressant effects.

We also examined whether the BLA network can support rhythmic oscillations. To the best of our knowledge, we are the first to show that the mouse BLA can generate *ex-vivo* oscillations at gamma-band range^**~40Hz**^ during high K^+^/Kainate (K^+^/KA) tone (Fig. 4, Supplementary Fig. 3). Gamma power is stronger in the BLA than nearby regions in awake mice, and BLA multi-units are phase-locked to BLA gamma oscillations(Kanta, Pare, and Headley 2019). Together, these findings indicate that the BLA can generate and self-sustain network oscillations in the gamma-band range despite the lack of an apparent laminar organization with parallel pyramidal cell organization. The generation of gamma rhythmogenesis in the *ex-vivo* BLA seems to rely exclusively on GABAergic interneuron signaling as these gamma-oscillations were only blocked by a GABA_a_R blocker and not by an AMPAR blocker (Fig. 4d-g), in agreement to gamma oscillations in the rat BLA(Randall, Whittington, and Cunningham 2011). Furthermore, prolonged exposure of a GABAA receptor antagonist induced oscillations in the theta band-range^**3-12Hz**^ (Fig. 4f-g). Therefore, it is possible that under certain conditions, such as reduced inhibitory tone, the BLA can generate theta-oscillations intrinsically that can be detected by LFP recordings.

We also demonstrate that rhythmic photo-excitation of PV^+^ interneurons could entrain network activity in BLA slices (Fig. 5e-f, Supplementary Fig. 4b,c). PV^+^ interneurons form a large interconnected network in the BLA(Muller, Mascagni, and McDonald 2005) and release GABA onto the perisomatic regions of their post-synaptic neurons, allowing them to effectively control their output(Bartos, Vida, and Jonas 2007; Veres, Nagy, and Hájos 2017). Indeed, it has been shown that rhythmic optogenetic BLA PV+ activation in awake mice entrains BLA multi-unit activity(Ozawa et al. 2020). Thus, our findings together with existing literature suggest that PV^+^ interneurons are ideally poised to synchronize and control the oscillatory state of the BLA network and pharmacological agents targeting this cellular mechanism are capable of altering network and behavioral states.

We further observed that allo application enhances tonic inhibition on BLA PV^+^ interneurons but not on principal cells (Fig. 3), similar to previous reports(Liu et al. 2014). Tonic inhibition on interneurons has been previously proposed to promote network synchronization(Pavlov et al. 2014). It is possible that the ability of allo to modulate BLA network synchronization involves the potentiation of tonic inhibition on PV+ interneurons.

The current study focused on the impact of allopregnanolone on oscillations within the BLA. Future studies are required to resolve the impact of allopregnanolone on coupling between anxiety-related networks, such as communication between the BLA and medial prefrontal cortex or the ventral hippocampus(Calhoon and Tye 2015; Likhtik et al. 2014; Tovote, Fadok, and Lüthi 2015). Further, the impact of the saline injections on oscillations in the BLA, which interestingly oppose the effects of allopregnanolone (Fig. 2c & Supplementary Fig. 2a-b), are likely the result of the stress of the injection and suggest that acute stress may also impact network and behavioral states.

It was observed that the long-lasting antidepressant effect in clinical trials which outlasts the pharmacokinetics of the drug exposure and is not easily explained by the known mechanism of action (Daly et al. 2018; Meltzer-Brody et al. 2018). Our data demonstrate that allopregnanolone is able to shift the network oscillation state in the BLA through δ subunitcontaining GABAARs, though not exclusively through this mechanism (Fig. 2, Fig. 7j). One could envision that allopregnanolone acts on δ subunit-containing GABA_a_Rs to shift the network to a healthy network state that is more stable and can persist in the absence of the compound. Future studies are required to fully understand the long-term effects of allopregnanolone on the network, which will be informative for understanding the mechanisms mediating the clinical effectiveness of allopregnanolone as an antidepressant treatment. These data demonstrate a novel molecular and cellular mechanism orchestrating network and behavioral states, although, we do not presume that this is the only mechanism involved in switching between network states. In fact, there are likely numerous mechanisms which can alter the network in a similar manner(Swensen and Marder 2000).

## Author Contributions

Conceptualization & Writing-Original: P.A and J.L.M., Project Administration, Supervision & Funding Acquisition: J.L.M., Visualization: P.A and P.L.W.C, Data curation, programming and mouse LFP analysis: P.A., Writing-Review & Editing: P.A, J.L.M, P.L.W.C, N.L.W, G.L.W, A.C.S, D.P.N, M.K. and M.C.Q.

J.L.M conducted and analyzed whole-cell, mouse awake LFP and behavior experiments, P.A and P.L.W.C conducted ex-vivo LFP experiments, A.C.S, D.P.N, M.K. and M.C.Q conducted and analyzed Human and Rat EEG experiments. L.C.M conducted and analyzed quantification delta-subunit GABAA receptor experiments. N.L.W conducted the fMRI experiments. N.L.W and G.L.W analyzed the fMRI experiments.

## Acknowledgements

The authors would like to thank Dr. Leon Reijmers for providing constructive comments on the manuscript. The authors would also like to thank the Biostatistics, Epidemiology, and Research Design (BERD) Center and specifically Dr. Jessica Paulus, ScD, for consultation and assistance with the correlational analysis and statistical measures. Collectively we also thank the Center for Translational Neuroimaging at Northeastern University for providing access to the Bruker BioSpec 7.0T and for assisting in fMRI data collection and analysis. J.L.M., P.A., N.L.W, G.L.W, P.L.W.C., and L.C.M. were supported by R01AA026256, R01NS105628, R01NS102937, and a sponsored research agreement with SAGE Therapeutics.

---

**Supplementary Figure 1 (Supporting Figure 1).**
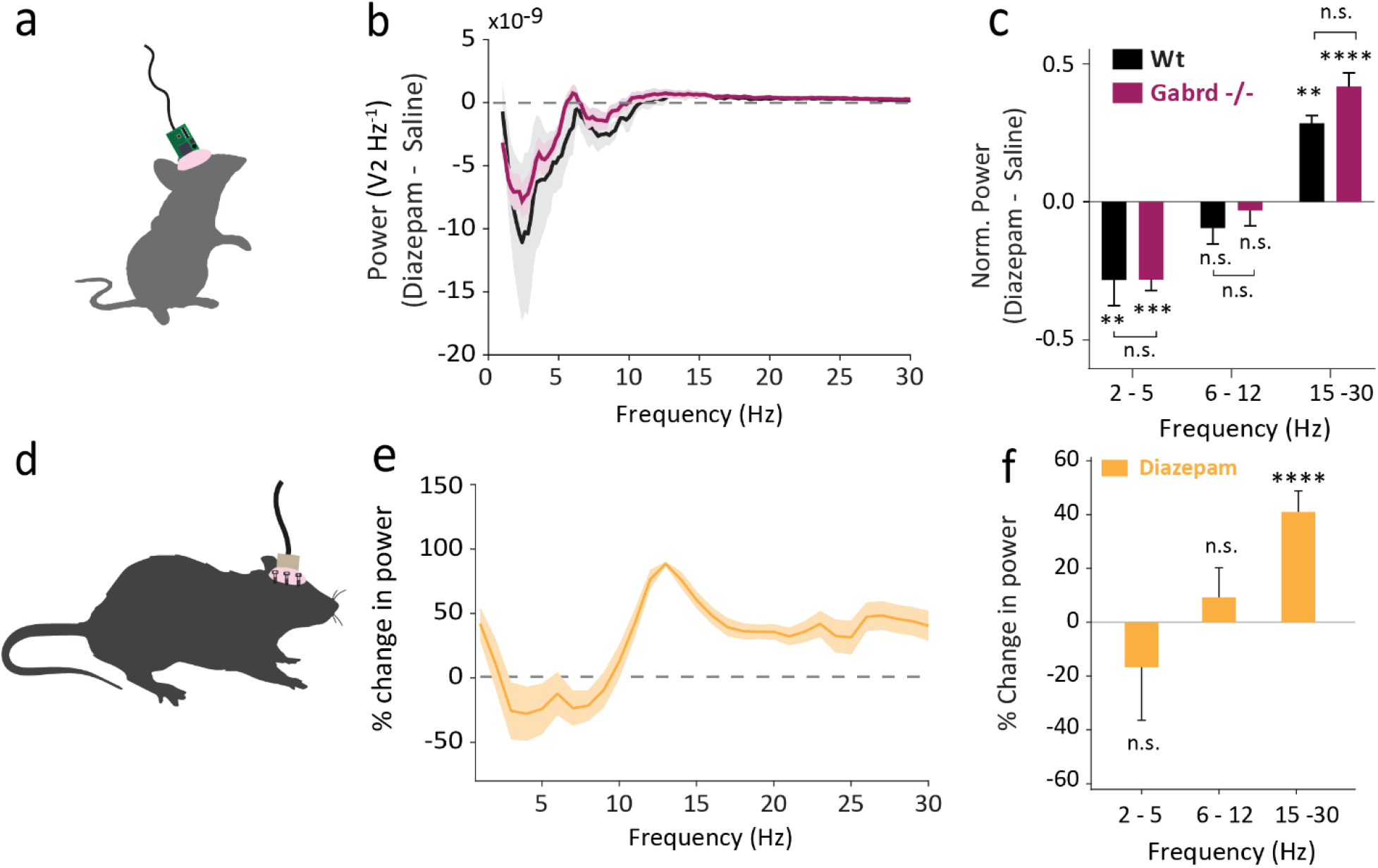
Diazepam treatment altered theta oscillations differentially from Neurosteroids. **a-c, mice; d-f, rats. a**, Schematic for BLA LFP recordings in awake mice. **b**, Power spectral density difference between diazepam (1 mg/Kg) and saline IP applications. **c**, Normalized power area difference between acute diazepam and saline treatment across multiple frequency bands; Single stars represent paired t-tests between drug treatments. Brackets represent unpaired t-tests between genotypes (n_Wt_=5 mice, n_Gabrd-/-_=5 mice). **d**, Schematic for EEG recordings in awake rats. **e**, Power spectral mean difference from vehicle in EEG power (defined as percentage change from baseline) in the rat with diazepam (orange). **f**, Mean (± SEM) change in power across frequency bands. Shaded regions and error bars represent SEM (n_diazepam_=18 rats); stars represent unpaired t-tests.

**Supplementary Figure 2 (Supporting Figure 2).**
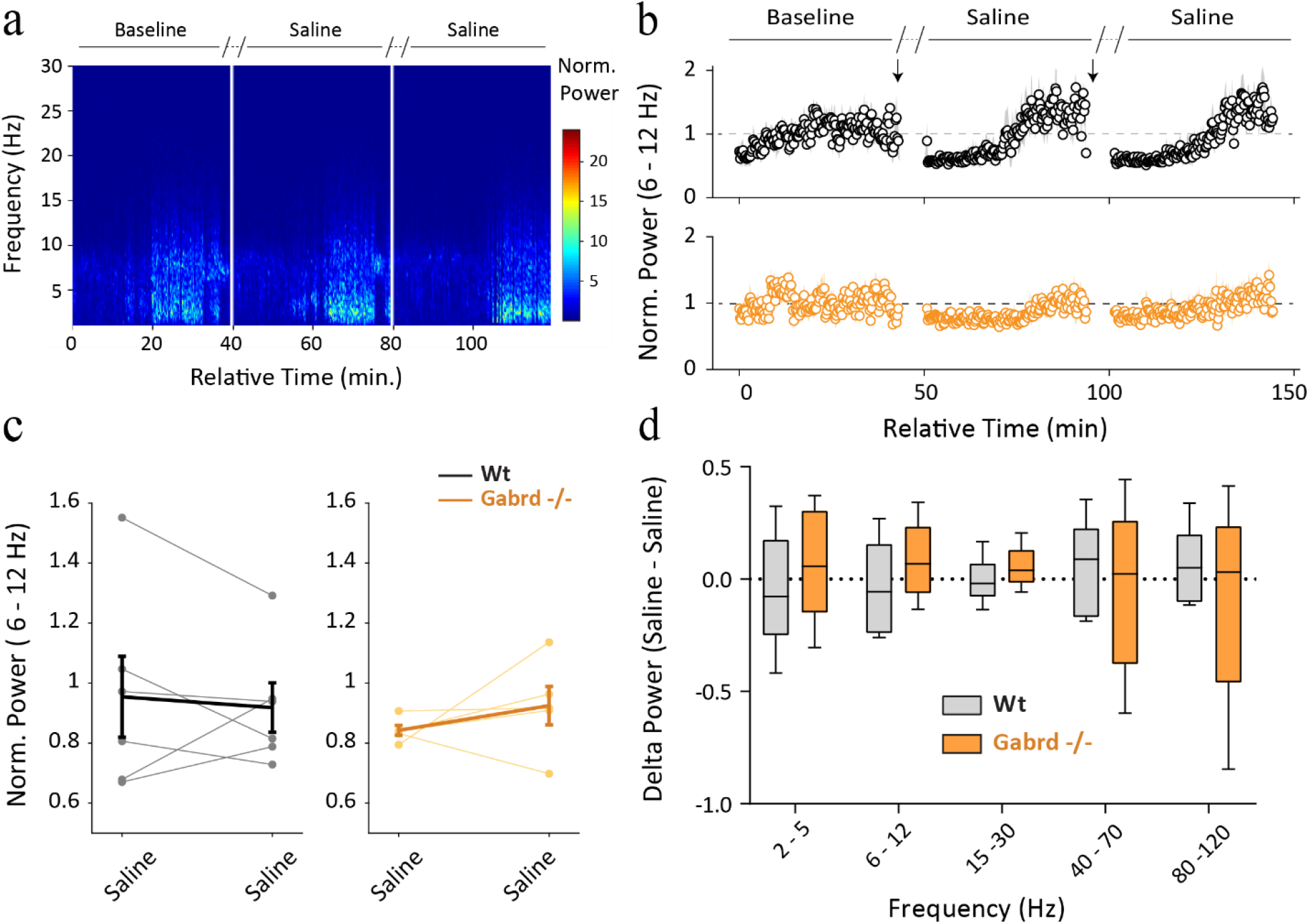
Repeated IP saline application did not alter BLA network oscillations oscillation. **a**, Representative spectrogram of BLA oscillations during repeated IP saline application. Power area^**6-12Hz**^ normalized to baseline during repeated IP saline application **b**, Across time **c**, Average dot plot; light color lines indicate individual experiments, dark lines the average and error bars the SEM. **d**, Normalized power area difference between repeated saline application across multiple frequency bands. Two-way repeated measures ANOVA was used to assess the interactions between frequency bands and repeated saline injections. Wt: F(4,25)=0.45, p=0.77; Gabrd F (4, 20) = 0.30, p=0.87). n_Wt_=6 animals, n_Gabrd-/-_=5 animals.

**Supplementary Figure 3 (Supporting Figure 4).**
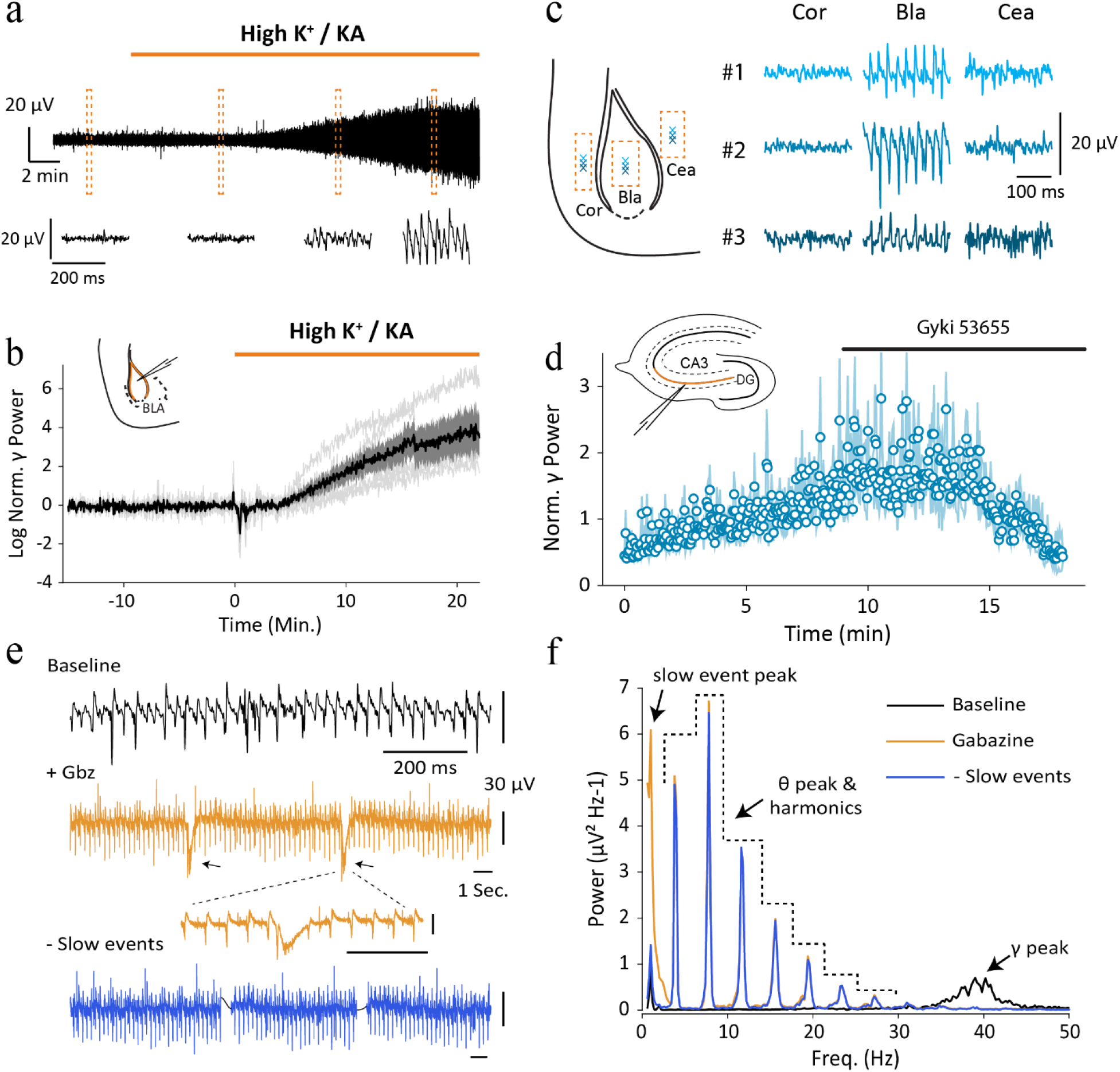
**a-b**, Application of kainic acid (800 nM) in aCSF with elevated KCl (7.5 mM) induces gamma oscillations. **a**, Top-Representative LFP trace and Bottom-Magnified traces from orange rectangles. **b**, Normalized gamma peak power^**30-80Hz**^. Grey traces represent individual experiments (n=9 slices), black trace the mean and shaded area the SEM. **c**, Gamma oscillations can only be recorded in BLA and not surrounding areas; Left-schematic of approximate LFP recording positions in Right-rows represent LFP recordings from separate slices (n=3 slices). **d**, Gyki 53655 application (20 μM) suppresses carbachol-induced (5 μM) oscillations in CA3; dots represent mean and shaded region represents SEM. **e-f**, Induction of additional slow events from gabazine application in BLA. **e**, Representative LFP recordings from BLA and **f**, respective power spectral density.

**Supplementary Figure 4 (Supporting Figure 5).**
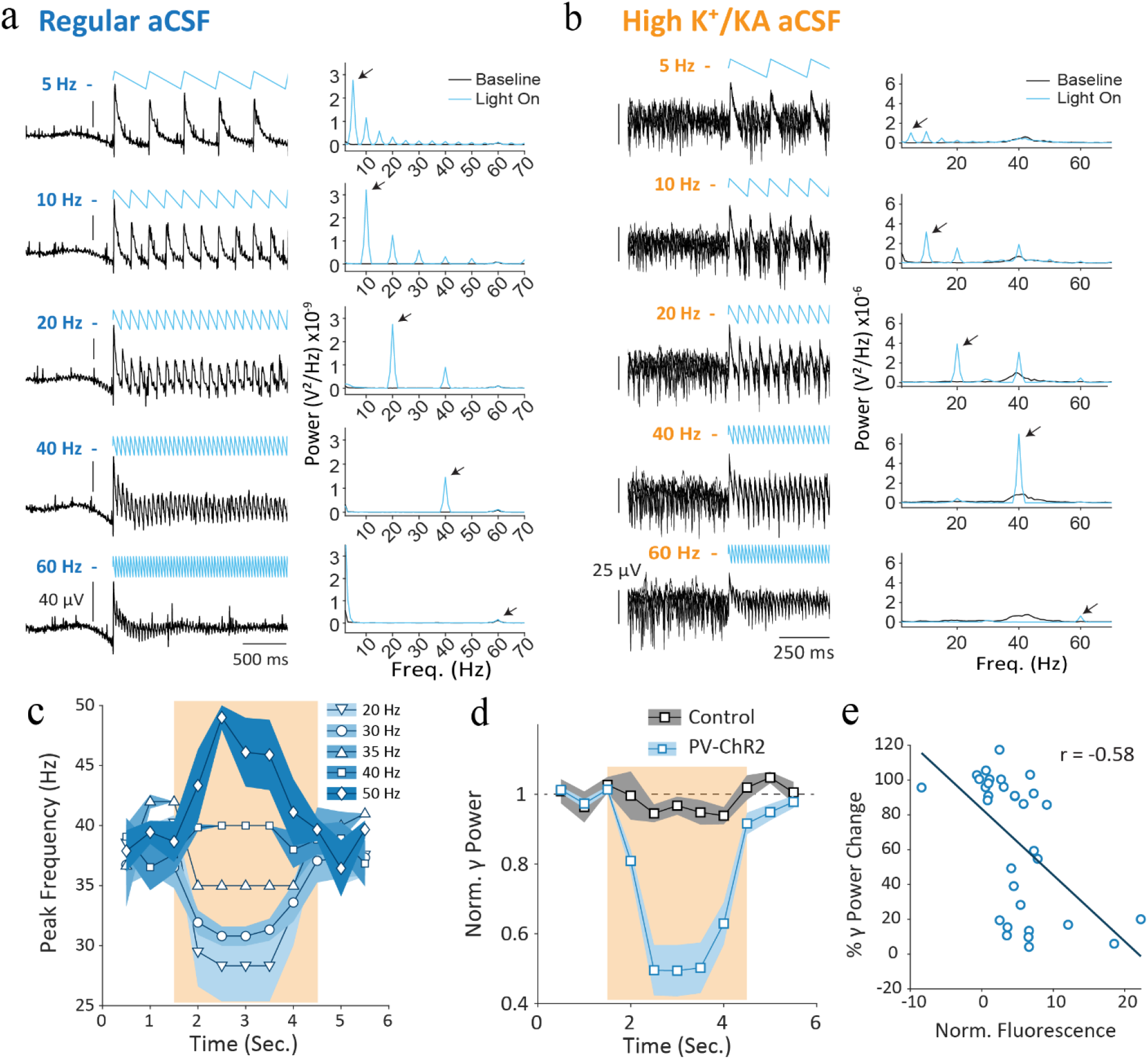
**a-b**, Left-Representative LFP recordings from BLA and Right-associated PSDs obtained during PV^+^ interneuron pulsed photo-excitation (black arrows indicate peaks at stimulation frequency). **c**, Average Peak frequency of oscillations during PV^+^ interneuron pulsed photo-excitation, shaded region=SEM; n slices: 20Hz=12, 30Hz=15, 35Hz=3, 40Hz=13, 50Hz=9. **d**, Average Normalized gamma power to baseline in controls (n=6 slices) and PV-ChR2 animals during sustained photo-excitation, shaded region=SEM (n=23 slices). **e**, Gamma Power change during sustained PV^+^ interneuron photo-excitation plotted against normalized BLA fluorescence (r=-0.58, p=0.0005, n=32 slices, Spearman Correlation). Light intensity at 10 mW.

**Supplementary Figure 5 (Supporting Figure 6).**
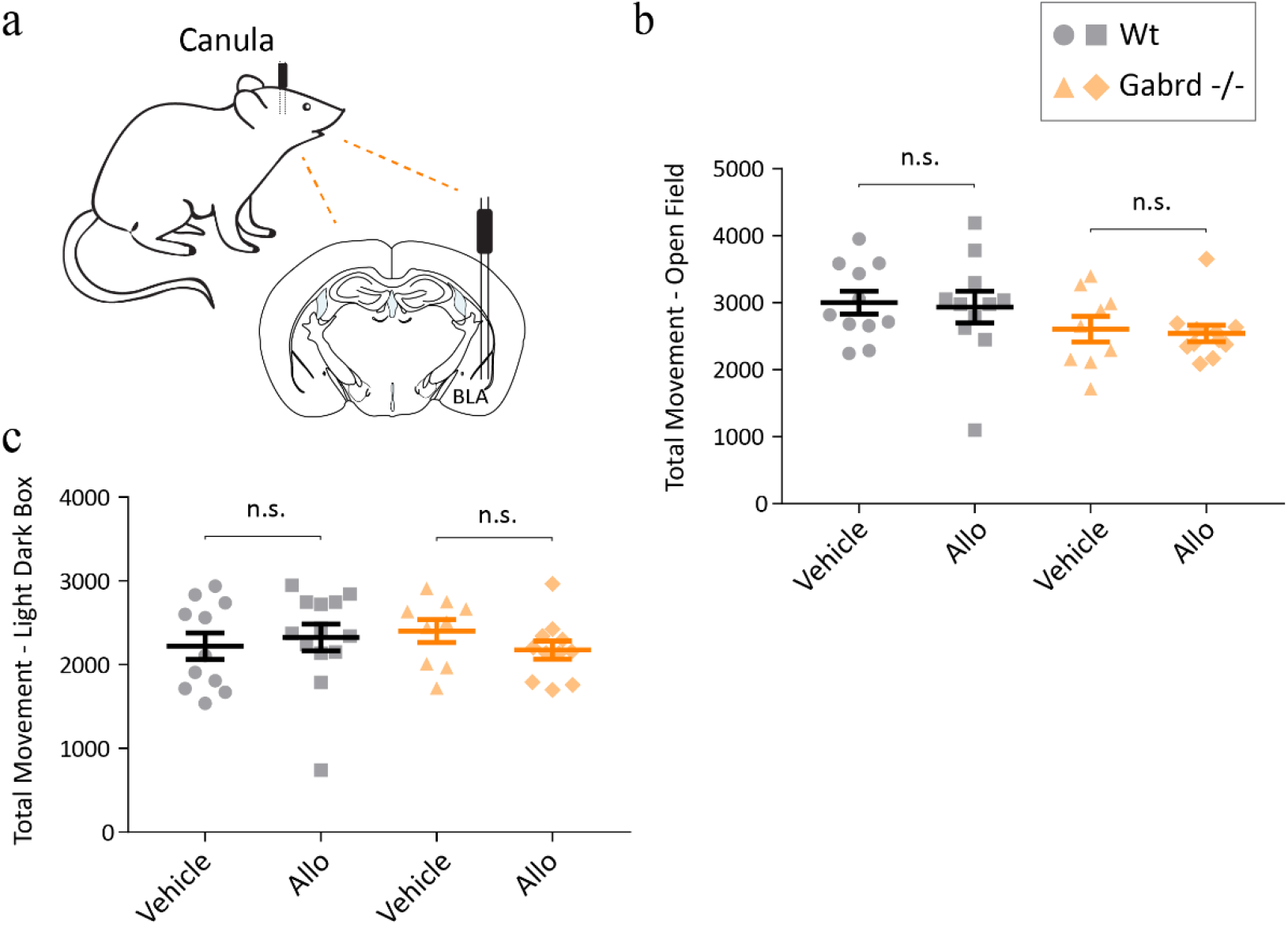
**a**, Schematic for canula implantation in basolateral amygdala (BLA). Total movement (# total beam breaks) in **b**, Open Field **c**, Light Dark Box. Error bars represent SEM. Brackets represent unpaired t-tests between vehicle against allo treated groups.

