## Supplement 1 for "Allopregnanolone mediates affective switching through modulation of oscillatory states in the basolateral amygdala"

### Supplemental Methods

#### Animals

Adult mice (>P60) were housed at Tufts University School of Medicine in a temperature and humidity-controlled environment. All animal procedures were handled according to the protocols approved by the Tufts University Institutional Animal Care and Use Committee (IACUC). Animals used include: C57BL/6J, global Gabrd$^{-/-}$(Mihalek et al. 1999), PV-Gabrd$^{-/-}$(Lee and Maguire 2013), PV-cre (B6;129P2-Pvalbtm1(cre)Arbr/J)(Hippenmeyer et al. 2005).

#### Stereotaxic Surgery

Mice were anesthetized by intraperitoneal injection (IP) of a ketamine/xylazine cocktail (90 – 120 mg/Kg and 5 – 10 mg/Kg, respectively). Before the onset of the procedure, sustained release-buprenorphine (0.5 – 1 mg/Kg) was administered subcutaneously as post-operative analgesia. The head of the mouse was shaved, antiseptic solution was applied, and an incision on the scalp was made to expose the skull. For viral injections, bilateral burr holes were made in the skull (From Bregma: −1.45 mm AP, ± 3.3 mm ML, −4.5 mm DV) and mice were bilaterally injected with 300-400 nL viral titer of AAV8-EF1a-DIO-hChR2(H134R)-mCherry-WPRE-HGHpA (#20297, Addgene, Karl Deisseroth) using a 33-gauge Hamilton syringe (infusion rate of 100 nL/min) in the BLA and the wound was closed using sutures. Animals were transferred to a temperature-controlled chamber for post-operational care until awake and were then monitored until recovery.

#### Behavioral Testing

C57BL/6J and global Gabrd$^{-/-}$ mice were implanted with a canula (Plastics One) in the BLA (AP - 1.35 mm, ML ± 3.45 mm, DV - 5.15 mm). Following a 3-week recovery, mice were infused either with saline solution (0.9 % NaCl), vehicle (15% 2-hydroxypropyl-β-hyclodextrin (HBPCD)), or 5 µg Allopregnanolone (2.5 µg/µl, Tocris) 30 minutes prior to behavioral testing.

Open-Field: The open field test was performed as previously described(Lee, Sarkar, and Maguire 2014; Sarkar et al. 2011). Mice were placed in the center of a 40 cm × 40 cm open field photobeam frame with 16 × 16 equally spaced photocells (Hamilton-Kinder) and the movement throughout the apparatus was measured over a 10-minute period. The number of entries, the total distance traveled, the amount of time spent in the center of the open field, as well as the total number of beam breaks were measured using an automated MotorMonitor software (Hamilton-Kinder).

Elevated-Plus Maze: The elevated plus maze was performed as previously described(Melón et al. 2019). Mice were place individually into the center of the elevated plus maze, elevated 75cm above with ground with two opposing 38cm x 6.5cm wide arms, one enclosed and one exposed. The closed arms have 10cm high walls and all four arms have 48 equally spaced photocells. The number of entries, distance traveled, and total time spent in the open arm was measured during the 10-minute test using an automated software (Ethovision).

Light-Dark Box: This light/dark box test was performed as previously described(Melón et al. 2019; Sarkar et al. 2011). Mice were individually placed in the dark compartment of a two chambered light/dark apparatus enclosed by a 22 cm x 43 cm photobeam frame with 8 equally spaced photocells (Hamilton-Kinder). The number of entries, distance traveled, and amount of time spent in the light compartment was measured using automated MotorMonitor software (Hamilton-Kinder) over the 10-minute test.

Tail Suspension: Mice were individually suspended by the tip of the tail from a bar at a height of 36 cm above a table for a 6 min test as previously described(Lee, Sarkar, and Maguire 2014). Each trial was videotaped and subsequently scored for the latency to the first bout of immobility and the cumulative time spent immobile. Mice were excluded from analysis if they climbed their tail or freed themselves during testing.

#### LFP recordings in awake mice

LFP recordings were performed in awake C57BL/6J and global Gabrd$^{-/-}$ mice, using in-house modified prefabricated headmounts (Pinnacle 8201). The headmounts were mounted to the skull using stainless steel screws as ground, reference, and frontal cortex EEG (AP +0.75 mm, ML ± 0.3 mm, DV −2.1 mm) electrodes. One depth electrode (PFA-coated stainless-steel wire, A-M systems) was placed into the BLA (AP −1.35 mm, ML ± 3.45 mm, DV −5.15 mm) and was used to record the LFP. Animals were tethered to the apparatus and the LFP was acquired at 4 KHz and amplified x100. The LFP data were band-pass filtered (1-300 Hz, Chebyshev Type II filter) and spectral analysis was performed in MATLAB using custom-made scripts utilizing the fast Fourier transform(Frigo and Johnson 2005). Briefly, recordings were divided into 5 second overlapping segments (50% overlap) and the power spectral density for positive frequencies was obtained by applying a Hann window to eliminate spectral leakage. Subsequently, data were merged to 30 or 300 second bins, the mains noise was eliminated from each bin between the 58-60 Hz band and replaced using the PCHIP method (piecewise cubic hermite interpolating polynomial). Values that were 5x larger or smaller than the median were removed and replaced with the nearest bin. For acute experiments, a baseline period was recorded for each animal followed by an IP injection with saline (0.9% NaCl) and a subsequent IP injection with the drug of interest (Allopregnanolone: 10 mg/Kg, SAGE-516: 5 mg/Kg, diazepam: 1 mg/Kg). The baseline period was used to normalize the power for each animal.

#### LFP recordings in *ex-vivo* brain slices

Mice were anesthetized with isoflurane and decapitated with a guillotine. Brains were extracted in ice cold (0-4$^oC$) sucrose solution containing (in mM) 150 sucrose, 15 glucose, 33 NaCl, 25 $NaHCO_3$, 2.5 KCl, 1.25 $NaH_2P0_4$, 1 $CaCl_2$, 7 $MgCl_2$ (300 – 310 mOsm). Coronal brain slices (350 µm) were obtained using the Leica VT1000s Vibratome and slices were transferred to an interface storing chamber in a warm (~ 34$^oC$) oxygenated aCSF solution containing (in mM) 126 NaCl, 10 glucose, 2 $MgCl_2$, 2 $CaCl_2$, 2.5 KCl, 1.25 $NaHCO_3$, 1.5 Na-pyruvate, 1 L-glutamine (300 – 310 mOsm) (normal). The hippocampus was dissected away to ensure that no hippocampal network activity propagated to BLA. All solutions were continuously bubbled with 95% $O_2$ and 5% $CO_2$. Slices were left to recover for at least 1 hour and were then transferred to an interface recording chamber where the local field potential (LFP) was obtained by inserting a borosilicate glass electrode into the BLA. Data were acquired through LabChart (ADInstruments) at 10 KHz and were low pass filtered at 3 KHz during acquisition. Spike artifacts were minimized using an amplitude cut-off (50 µV), the LFP data were band-pass filtered between 3-300 Hz (0.6-300 Hz for gabazine experiments) and spectral analysis was performed as before (refer to the LFP recordings in awake mice section). Gamma oscillations were induced by perfusing warm aCSF (~ 34$^{o}$C) with elevated potassium (7.5 mM KCl) and 800 nM Kainic acid. For removal of slow events in gabazine experiments, during analysis, data were low-pass filtered at 3 Hz and events were detected using a 20 µV threshold. Slow events were removed and replaced using the PCHIP method in the surrounding 600 ms segment. For ChR2 experiments, blue light was delivered through a fiber optic (200 µm, 0.22 NA) coupled to a blue laser (473 nm, max power=500 mW, Laserglow technologies). Waveforms for optical stimulation were created using a function generator (Agilent – 33210A) and were controlled by an Arduino Uno. The power spectral densities for optogenetic experiments were obtained in 1 second bins (50 % overlap) using custom-made MATLAB scripts. For LFP power ratio quantification the power of the oscillations was obtained around +/- 2 Hz of each stimulation frequency during (0 − 2 sec.) and before stimulation (0 − -2 sec.). The LFP power ratio was quantified by dividing the power during the stimulation period by the power before stimulation. The Continuous Morse Wavelet Transform(Lilly and Olhede 2012) was used to visualize opto-spectrograms.

Viral Expression: Brain slices were fixed in 4% paraformaldehyde (PFA) overnight and were subsequently washed with phosphate buffered saline (PBS) and kept at 4°C for short-term storage. Slices were then mounted on glass slides using VECTASHIELD® Hardset™ mounting medium and imaged using a digital epifluorescence microscope (Keyence). For optogenetic experiments, images were acquired using a 2X objective (0.1 NA). Slices with undetectable mcherry-fluorescence expression were excluded from further analysis. For correlation of gamma power change to BLA fluorescence, all slices were included, and their images were converted to

16-Bit using the Fiji software package. The fluorescence levels of for each slice was quantified using the difference between the mean of the whole slice subtracted from BLA.

#### Voltage-Clamp Recordings

Brains were extracted in ice cold (0-4$^{o}$C) aCSF solution containing (in mM) 126 NaCl, 10 glucose, 2 $MgCl_2$, 2 $CaCl_2$, 2.5 KCl, 1.25 $NaHCO_3$, 1.5 Na-pyruvate, 1 L-glutamine (300 – 310 mOsm) and 3 mM kynurenic acid. Coronal brain slices (350 µm) were obtained and transferred to a submerged chamber in a warm (~ 34$^{o}$C) normal aCSF for at least 1 hour prior to recordings. Slices were then transferred to a submerged recording chamber. Borosilicate glass electrodes were pulled with a resistance of ~3-5 MΩ (DMZ Universal Puller). The intracellular recording solution contained (in mM) 140 CsCl, 1 $MgCl_2$, 10 HEPES, 4 NaCl, 0.1 EGTA, 2 Mg-ATP, and 0.3 Na-GTP (pH=7.25, 280–290 mOsM). Data were acquired at 10 KHz and were low pass filtered at 3 KHz (AD Instruments). Recordings were performed in principal neurons in the BLA identified based on morphology and in visually-identified $PV^+$ interneurons using a PV-cre-AI9 mouse that labels $PV^+$ cells with td-Tomato fluorophore. To unmask the contribution of tonic current > 200 µM of Gabazine were perfused. The magnitude of the tonic-current was calculated as previously described (J. L. Maguire et al. 2005). Briefly, the mean current was measured during 10 ms epochs collected every 100 ms throughout the recording. A histogram of the baseline holding current (1 min prior to the addition of Gabazine) and in the presence of Gabazine (1 min after Gabazine blockade) were fit with a Gaussian and the difference in the mean of the fitted curve was attributed to the tonic current. Cells were excluded from analysis if series resistance or whole cell capacitance changed > 20% during the course of the recording.

#### EEG recordings in awake rats

Studies were conducted in line with the Guide for the Care and Use of Laboratory Animals, eighth edition (2011)(National Research Council (US) Committee for the Update of the Guide for the Care and Use of Laboratory Animals. 2011), US Public Health Service Policy on the Humane Care and Use of Laboratory Animals(PHS Policy on Humane Care and Use of Laboratory Animals | OLAW n.d.) and the Animal Welfare Act. Two rat studies were performed. For the first rat study, SGE-516 (dose levels of 1, 3, 10 and 30 mg/kg, n=6 rats per group) was formulated in 15% HBPCD and delivered intraperitoneally. In a second rat study, SAGE -217 (dose levels of 0.3, 3, 20 mg/kg, n=6 rats per group) was formulated in 30% sulphobutylether-cyclodextrin (SBECD) vehicle and delivered by mouth. After a 7-day acclimation period, Male Sprague-Dawley rats were anesthetized with either 70 mg/kg pentobarbital sodium (i.p., SAGE-217) or 30 mg/kg Zoletil and 3 mg/kg Xylazine mixture (0.6 ml/kg, SAGE-516) and implanted with three skull screws (negative screw: 2 mm anterior to bregma, 1.5 mm left; positive screw: 5 mm posterior to bregma, and 2.5 mm right; and reference screw: over cerebellum). After full recovery, rats were randomly assigned to a dose group. Before each dosing session, a 1-hour continuous baseline EEG was recorded, followed by dosing, and a 6-hour continuous EEG recording. EEG recordings were tethered, and EEG data were recorded using a A-M Model 1700 Differential Amplifier System at a sampling rate of 500 Hz per channel. Signal processing and statistics of the rat EEG were performed using MATLAB (Mathworks; Natick, MA). Pre-processing of the EEG involved artifact removal (deleting signal with amplitude greater than 5 times its standard deviation). The power spectrum was computed using FFT across time in sections of length 100, windowed with a

Hamming window, with 50 samples of overlap. EEG power density was computed from 1-50 Hz. In each of the two rat experiments multiple doses of drug were applied. To generate equivalent power spectra for comparison between drugs, the spectrogram for each animal over all doses was computed and segregated into 30-minute blocks. Blocks with power at 13 Hz in the 75-100% range were identified, and the average power spectra was computed in that block. For the SGE-516 data, 27 time points were located with appropriate power and averaged. Twenty similar time points were located in the SAGE-217 group. This method was selected as a means of generating snapshots in the power spectral density where, in the absence of pharmacokinetic data, the animals exhibited increased beta similar to that observed in the human EEG.

#### EEG recordings in human subjects

EEG was recorded using the Cadwell Easy II Amplifier (Kennewick, WA) and 19 electrode configuration according to the International 10-20 system (Fp1, Fp2, F7, F3, Fz, F4, F8, T3, C3, Cz, C4, T4, T5, P3, Pz, P4, T6, O1, O2, and reference to linked mastoids), under 5 minutes of eyes closed conditioned, with electrode impedance kept under 10kΩ. The times of recording were approximately 20 minutes before dosing and 2 hours post-dose. SAGE-217 was dosed at 30 mg in oral solution (40% HBPCD and 0.0025% sucralose) as part of a multiple-ascending dose (MAD study) in 7 healthy volunteers(Hoffmann et al. 2019). EEG were originally sampled at 2.4 kHz per channel using a -48dB anti-aliasing filter. Right before EEG processing, the EEG were band-pass filtered in the range of 0.53 Hz to 70 Hz. Pre-processing of the EEG included band pass filtering, gross voltage or EMG artifact rejection, bad channel interpolation, EOG artifact removal, head movement or other non-cerebral artifacts and cleaned data was then submitted to power spectral analysis (FFT with 1Hz resolution).

#### Quantification of GABAR-δ subunit containing cells

Parvalbumin specific-Gabrd knockout mice (PV-Gabrd$^{-/-}$) were generated by crossing floxed Gabrd mice(Lee, Sarkar, and Maguire 2014) with Parvalbumin-Cre mice obtained from Jackson Lab (Stock #012358). Genotyping for Parvalbumin-Cre was performed in house using the primers listed below to distinguish the Parvalbumin-Cre product. Genotyping for the floxed Gabrd was performed in-house with the primers listed below to distinguish the wildtype Gabrd (449 bp) or floxed-Gabrd (543 bp) product.

*Parvalbumin:*

5': CATGAGGAGTGGCATACACG

3': TTCGCGATTTGAGGTCTTCT

*Gabrd:*

5': GACTCCAGTTGCCAAGCCTTTAATTCC

3': CATCTGCCTGTACCTCCAATGCCTG

Immunohistochemistry was performed as previously described(J. Maguire and Mody 2009; Melón et al. 2019). Mice were anesthetized with isoflurane, euthanized by rapid decapitation and brains removed for overnight immersion fixation (4% PFA at 4°C). For δ-DAB, free-floating 40 µm coronal sections were quenched of endogenous peroxidase activity with 3% $H_2O_2$ in methanol for 30 minutes then incubated in 0.05 M citrate buffer at 90°C for 45 minutes, to improve antigen retrieval. Tissue was blocked in 10% normal goat serum in 0.3% PBS/triton for 1 hour and incubated with rabbit anti-δ-$GABA_AR$ (1:250,PhosphoSolutions 868A-GDN) overnight at 4°C. Following incubation with a biotinylated goat anti-rabbit secondary (Vector Laboratories, PK-6101), an avidin-biotin complex based detection system (Vector Laboratories, PK-6101) and DAB based HRP substrate (Vector Laboratories, SK-4100) was used to develop signal.

For parvalbumin and δ co-immunofluorescence, free floating 40 µm coronal sections were incubated in 0.5 M citrate buffer for 45 minutes at 90°C. Tissue was blocked in 10% normal goat serum in 0.3% PBS/triton for 1 hour and incubated with rabbit anti-δ-$GABA_AR$ (1:250,PhosphoSolutions 868A-GDN) and mouse anti-parvalbumin (1:1000, Sigma P3088) for 4 days at 4°C. Slices were washed and incubated with biotinylated goat anti-rabbit (1:1000, Vector Laboratories BA1000) and Alexa-Fluor 647 conjugated goat anti mouse (1:200,ThermoFisher Scientific A28181) for 2 hours. Slices were incubated for 2 hours with streptavidin conjugated-Alexa-Fluor 488 (1:200, ThermoFisher Scientific S32354) before being mounted on slides with antifade hardset mounting medium with DAPI (Vectashield H1500).

#### Chronic unpredictable stress (CUS) protocol

Adult male C57BL6/J mice from Jackson Laboratory were housed at Tufts University School of Medicine's Division of Laboratory Animal Medicine facility on a 12 h light/dark cycle (lights on at 0700 hrs). Mice were housed in clear plastic cages (4 mice per cage) with ad libitum access to food and water. All procedures were approved by the Tufts University Institutional Animal Care and Use Committee. For chronic unpredictable stress (CUS) mice underwent a three-week protocol consisting of alternating overnight stressors (unstable cage, overnight illumination, restricted food and water, restricted cage access, or cage tilt) for four nights, followed by a fifth consecutive day of an acute stressor (restraint or swim stress). Sage treated mice received the SGE-516 compound in their chow.

**Example CUS Paradigm (3 weeks - 5 days: Monday to Friday)**

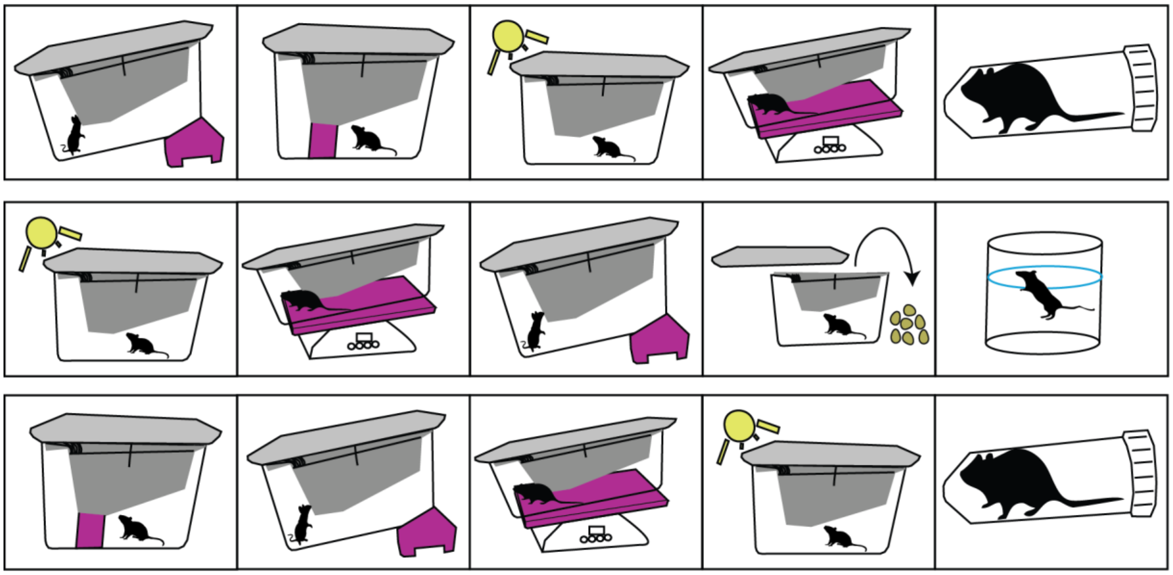

#### fMRI

Mice were acclimated to the imaging environment for 15, 20, 25, 30, and 35 minute increments over five consecutive days before imaging was conducted. During the acclimation period mice were anesthetized with isoflurane before being placed into a small rodent holder tube made for small rodent magnetic resonance imaging. Subsequent to recovering from anesthesia mice were placed in a dark box with a prerecorded audio of the MRI pulses.

CUS/Sage mice (n=5) underwent the 3-week chronic stress protocol while receiving the SGE-516 compound in their chow. Sage treated mice (n=5) received the SGE-516 compound in their chow and were gently handled by the experimenter twice per week. Control mice (n=5) were gently handled by the experimenter twice per week. All mice underwent the imaging acclimation protocol.

Magnetic Resonance Imaging was conducted using a Bruker BioSpec 7.0T with a 20-cm horizontal magnet and 20-G/cm magnetic field gradient quadrature transmit/receive coil (ID 38mm) at the Center for Translational Neuroimaging at Northeastern University. High-resolution anatomical scans were acquired by a Turbo Spine-Echo pulse sequence (0.75 mm; FOV 1.8 cm; data matrix 256 x 256; TR 2.1 sec; TE 12.4 msec; Effect TE 48msec; NEX 6) with an acquisition time of 6.5 minutes. A multi-slice T-2 weighted pulse sequence was used to acquire functional images. The entire imaging session was conducted in less than 30 minutes for each mouse. Mice were individually anesthetized at the beginning of the session and positioned inside the coil. Oxygen saturation, respiratory rate (brpm), and heart rate (bpm) for each mouse were monitored throughout the sessions using a rodent-specific MouseOx Plus sensor system (STARR Life Sciences Corp., Oakmont, PA).

Functional scans were segmented and labelled in reference to a 3D Allen Mouse Brain Atlas with ~138 brain regions. Segmentation, labelling, and preprocessing were performed using 3D Slicer, AFNI/SUMA and FSL, and MATALB, respectively. The BOLD signal for each voxel (0.00706mm3) was averaged across subjects and a voxel-based analysis was conducted to calculate the percentage of signal change between voxels. Student's t-test with a 95% confidence level, two-tailed distributions, with the assumption of a non-Gaussian distribution were performed on each voxel. BOLD signal changes were normalized in reference to the volume of individual brain regions using the equation: normalized voxels per region = (Number of activated voxels in a region x 100)/(Total number of voxels in a region). Changes in BOLD signal between experimental groups were measured via one-way ANOVA of each MRI pulse (7 seconds per pulse over 7 minutes). To quantify connectivity, the correlation matrix for the BOLD signal across brain areas was calculated (Pearson correlation) and transformed to z-scores. The resulting z-score for the percentage of BOLD signal change between voxels was transformed into a matrix where all values without statistical significance were set to zero. A baseline of 2% signal change was used to account for normal physiologic changes in blood oxygenation in the awake mouse brain.

The effects of CUS were normalized to non-stressed animals by calculating the difference between the fMRI z-matrices of stressed and non-stressed animals (Z-CUS – Z-non-stressed). The effects of the sage compound on animals that underwent CUS were normalized to non-stressed animals that received the sage compound by subtracting those z-matrices (Z-CUS-sage – Z-sage). Finally, the normalized CUS and CUS-sage matrices were compared to determine if the effects of CUS were either reversed or potentiated by the sage compound. Only differences larger than 1 standard deviation were reported. For connectivity plots, all connections between brain areas were plotted with line thickness equal to the calculated z-score and color-coded base on outcome (Fig. 7j): (Blue) Upregulated by CUS, no change with sage, (Black) Downregulated by CUS, no change with sage, (Green) Up/downregulated with CUS and reversed by sage, (Pink) Up/downregulated by CUS, further potentiated by sage. Plots generated with nxvis, networkx, and matplotlib Python packages.

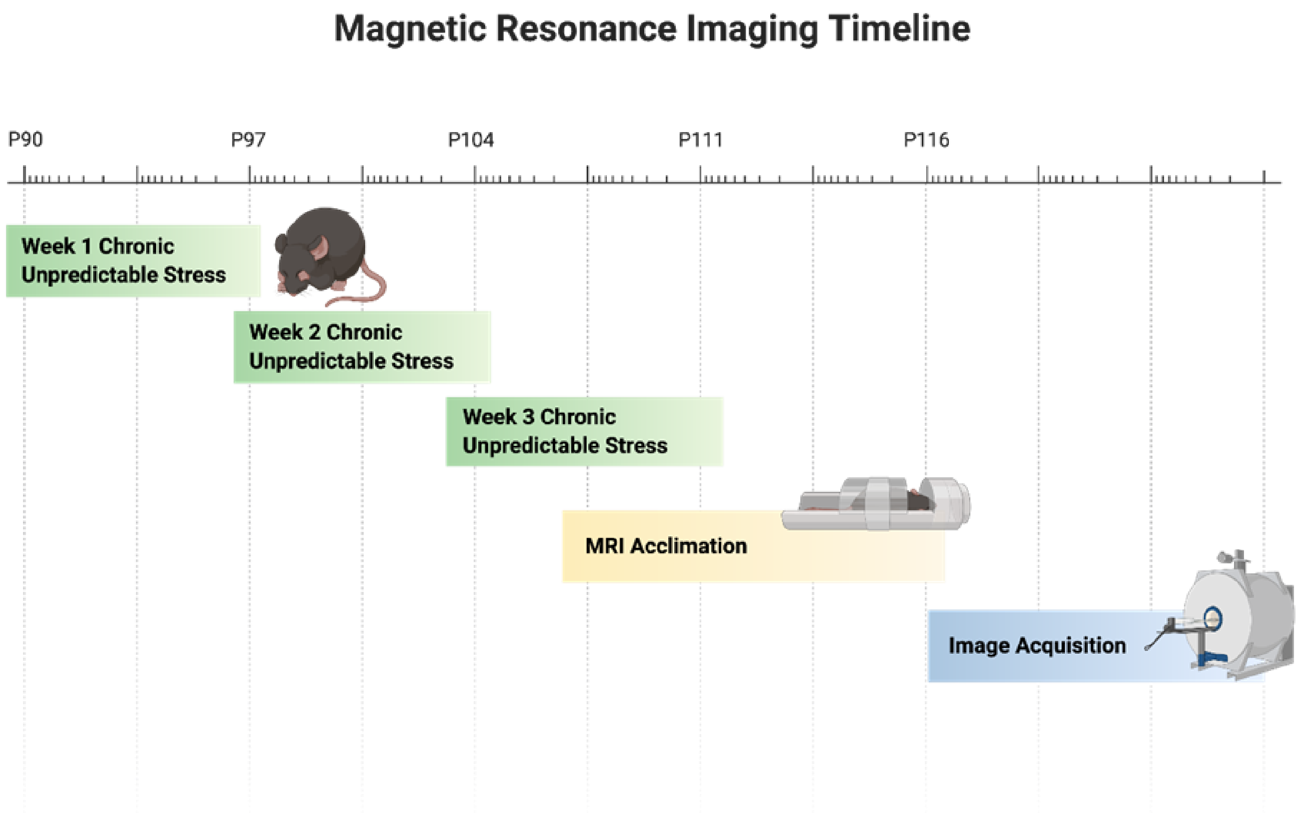


#### Quantification and statistical analysis

All statistical tests were performed using the PRISM 7 software (GraphPad) and MATLAB (Mathworks). All values were reported as mean ± SEM unless otherwise stated. P-values < than 0.05 were considered significant; $p \geq 0.05$=n.s. $p<0.05$=*, $p<0.01$=**, $p<0.001$=***, $p<0.0001$=****.

#### Data code and availability

The code for the MATLAB app for LFP and oscillation spectral analysis can be found on GitHub at https://github.com/pantelisantonoudiou/MatWAND. All other MATLAB/Python code used for data analysis will be provided upon reasonable request from the lead author Jamie Maguire. An interactive version of the fMRI connectivity plot (Fig. 7j) can be found at [http://fmri-circos.herokuapp.com](http://fmri-circos.herokuapp.com).
